# Kinetic Fingerprints as Mechanistic and Clinical Roadmaps Across KIT Activation States

**DOI:** 10.64898/2026.05.20.726488

**Authors:** Ana Corrionero, Niall Prendiville, Tatiana Cazorla, Maria Baena-Nuevo, Sandra Röhm, Emilio Camafeita, Stefan Knapp, Patricia Alfonso

## Abstract

In cancer therapy, traditional approaches often overlook the dynamic nature of drug-target interactions. We introduce kinetic fingerprints as a mechanistically informative tool to guide kinase inhibitor design and predict clinical performance. Profiling 172 compounds across multiple KIT conformations, including the oncogenic D816V mutation, show that prolonged residence time determines therapeutic success, while mutations accelerating dissociation rates (*k*_*off*_) drive resistance, positioning *k*_*off*_ as a robust predictor of clinical failure. Beyond efficacy and resistance, kinetic signatures map molecular behavior: fast-associating scaffolds engage readily populated KIT states, slow binders overcome conformational barriers like juxtamembrane repositioning, and extended residence times highlight ligands stabilizing regulatory elements (G-loop and regulatory spine). Kinetic profiling further unveils mechanisms invisible to conventional methods, such as drug-induced kinase degradation, and exposes selectivity dimensions beyond affinity: avapritinib exhibits durable KIT D816V engagement yet transient off-target binding. Our findings redefine the evaluation of KIT inhibitors, establishing a framework for rational, kinetics-guided drug discovery in KIT-driven cancers.

**Table of Contents:** Kinetic profiling of 172 compounds across KIT conformations—including D816V—reveals kinetic fingerprints that predict efficacy, selectivity, resistance, and inhibitors’ ability to stabilize key regulatory elements. Far from a secondary metric, binding kinetics provide mechanistic insights beyond affinity, offering a powerful framework for rational drug design in KIT-driven cancers.

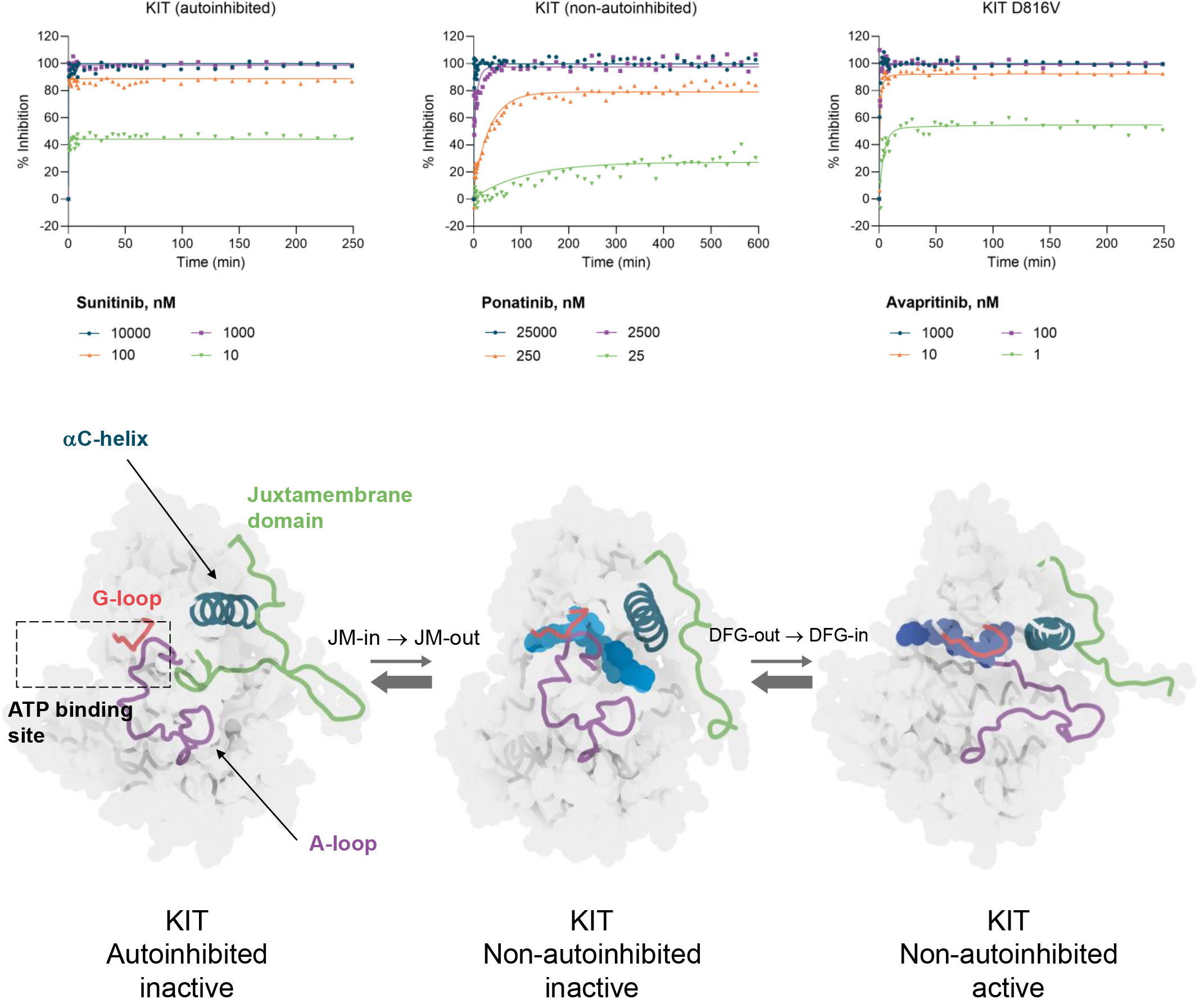

## Introduction

The KIT receptor, a member of the Type III receptor protein-tyrosine kinase family, which includes related proteins like PDGFRA, PDGFRB, FLT3 and CSF1R, plays a key role in cell growth, survival and differentiation by binding to stem cell factor (SCF).^1^ In the absence of SCF, KIT exists in an inactive autoinhibited state. The juxtamembrane (JM) domain forms a hairpin loop that inserts into the active site (JM-in), stabilizing the activation loop (A-loop) in a DFG-out conformation and maintaining the αC-helix in an inactive outward position (αC-helix out) (**Figure 1**).^2^

**Figure 1:**
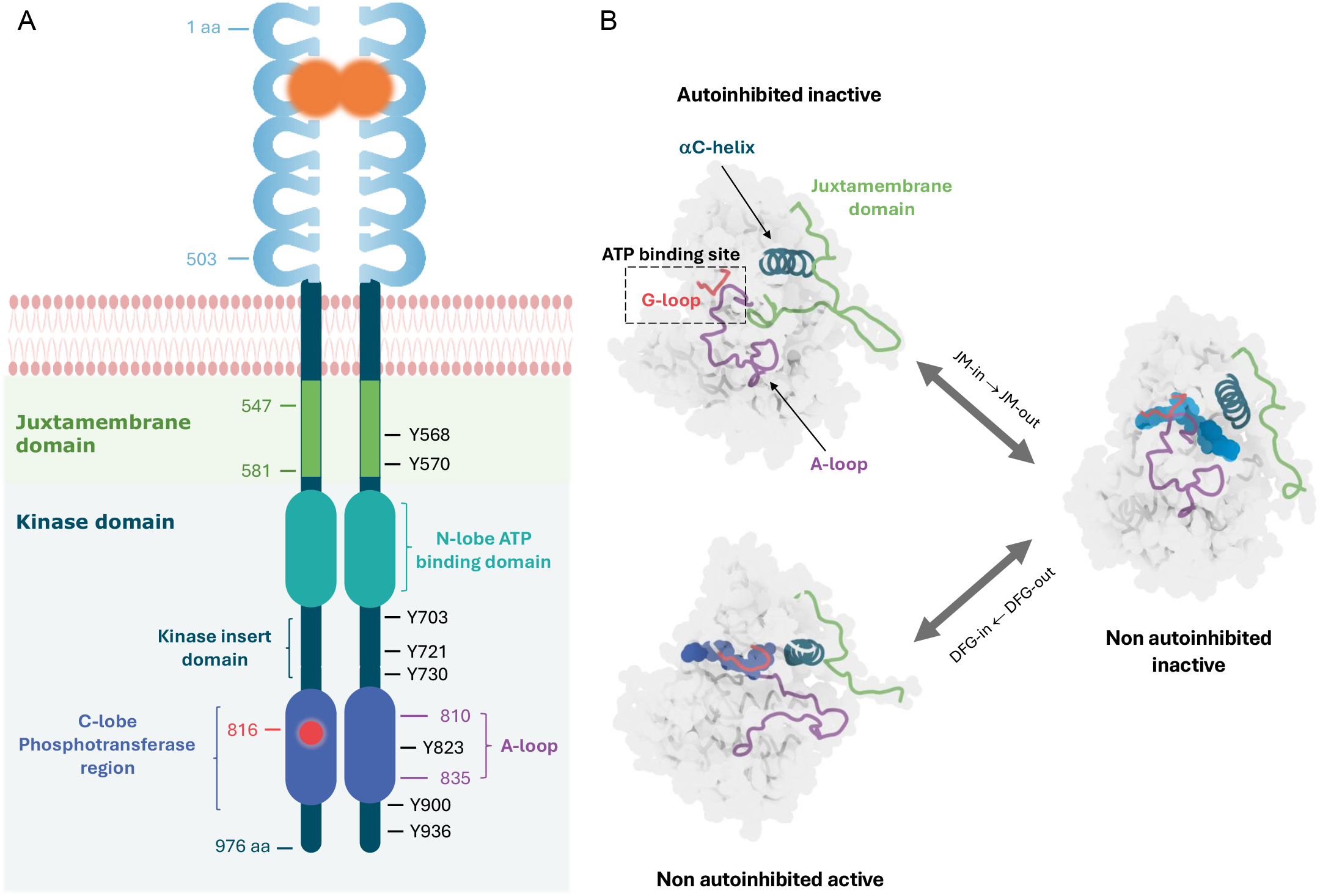
KIT receptor structure. (A) KIT spans 976 amino acids, with five extracellular immunoglobulin domains (23-520 aa) for SCF binding, a transmembrane helix (521-543 aa), a JM domain (544-581 aa) and a kinase domain (582-937 aa) with two lobes: a N-lobe containing the ATP-binding site and a C-lobe with the phosphotransferase region and A-loop (810-835 aa), separated by the Kinase Insert Domain (KID).^3^ (B) KIT receptor exists in at least three distinct activation states. The autoinhibited state, with the JM domain (green) blocking the active site and restricting A-loop (purple) activation (PDB: 4HVS). The inactive non-autoinhibited state with the phosphorylated JM domain displaced from the active site. In complex with the type II inhibitor ponatinib (blue, PDB: 4U0I). The active non-autoinhibited state stabilized in the DFG-in conformation and bound to the type I_1/2_ inhibitor avapritinib (violet, PDB: 8PQ9).

Upon SCF binding, KIT dimerizes which triggers transphosphorylation of four tyrosine residues in the JM domain, most notably Y568 and Y570. This phosphorylation displaces the JM domain into a solvent-exposed state (JM-out), releasing autoinhibition while the kinase remains inactive. Subsequent phosphorylation of Y823 within the A-loop stabilizes the active DFG-in conformation, accompanied by repositioning of the αC-helix (αC-helix in) and glycine-rich loop (G-loop), a flexible region that plays a crucial role in ATP binding. These conformational changes activate KIT, initiating downstream signaling pathways via phosphorylated tyrosines (Y547, Y553, Y568, Y570, Y703, Y721, Y730, Y823, Y900 and Y936) that serve as docking sites for SH2 domain-containing proteins (Figure 1).^3–5^

The constitutive activation of KIT is implicated in several malignancies, including gastrointestinal stromal tumors (GISTs), systemic mastocytosis (SM), acute myeloid leukemia and papillary renal carcinomas. Gain-of-function mutations disrupt KIT’s autoinhibitory regulation, driving uncontrolled signaling. In SM, over 90% of adult cases harbor the KIT D816V mutation, which causes partial unfolding of a short helix (residues 817–819) within the A-loop. This structural alteration disrupts JM domain dynamics and impairs the critical communication between these regions required to maintain the inactive state.^1,6^ This mutation also induces coordinated motions between the G-loop and αC-helix absent in KIT wild-type (WT), resulting in ligand-independent kinase activation and a ~586-fold increase in activity.^7^ Owing to its strong resistance to imatinib, the KIT D816V mutation requires alternative therapies; midostaurin and avapritinib are approved options for indolent and advanced SM due to their inhibitory activity against this variant.^2,4,9^

In nearly 80% of metastatic GISTs, primary activating mutations cluster in the JM domain or the extracellular domain of KIT.^1^ Though first-line imatinib therapy elicits early clinical responses, resistance often develops through secondary mutations, particularly in the ATP-binding pocket or A-loop. This has led to the development of subsequent therapies, including sunitinib, regorafenib, and ripretinib.^7,8^ Additionally, KIT overexpression occurs in malignancies such as chromophobe renal cell carcinoma, renal oncocytoma, colorectal cancer and small cell lung cancer.^6^ As a result, there is a sustained effort to develop effective and selective KIT inhibitors to address the challenges posed by resistance mutations and KIT overexpression.^10,13^

Although equilibrium-based affinity metrics (*K*_*d*_ or IC_50_) remain central to early drug discovery, they provide only a static snapshot of interactions that are intrinsically dynamic. As a result, they often fail to explain why compounds with similar affinities can elicit different pharmacological responses. Binding kinetics addresses this limitation by quantifying the association rate (*k*_*on*_), which reflects how quickly a drug engages its target, and the dissociation rate (*k*_*off*_), which determines how long a drug remains bound. These parameters often play an important role in anticipating both efficacy and selectivity. Drugs with longer residence times (τ), defined as 1/*k*_*off*_, frequently induce conformational shifts in their targets, stabilizing unique binding sites or activating specific states that enhance their therapeutic effects. Prominent therapeutic modalities where residence time impacts effectiveness include PROTACs, molecular glues and covalent drugs.^14–23^

Furthermore, extending residence time also helps counteract resistance caused by mutations that accelerate drug dissociation, thereby improving clinical outcomes. Finally, the interplay between residence time and pharmacokinetics further underscores its therapeutic relevance. While clearance determines systemic drug exposure, prolonged residence can extend target engagement beyond elimination and may limit metabolic degradation, ultimately influencing overall clearance.^24–28^

By integrating kinetic profiling into the analysis of 172 compounds across distinct conformational states of KIT— including its autoinhibited, non-autoinhibited and oncogenic D816V forms—this study reveals kinetic fingerprints that uncover mechanistic insights inaccessible to affinity-based approaches. These kinetic signatures provide valuable predictors of efficacy, selectivity, resistance profiles and inhibitors’ ability to stabilize key regulatory elements. Far from being a secondary parameter, binding kinetics emerge as a mechanistically rich layer of information that complements and extends affinity metrics. Taken together, this kinetic perspective establishes a powerful, mechanism-informed framework for the rational design of next-generation therapies in KIT-driven malignancies.

## Results and Discussion

### Large-Scale Binding Kinetics Profiling of Kinase Inhibitors Across KIT Activation States

To assess how activation state influences inhibitor affinity and binding kinetics, we used our KINETICfinder^®^ platform^29^ to develop high-throughput TR-FRET binding kinetic assays for the autoinhibited and non-autoinhibited states of KIT WT receptor, as well as the constitutively active KIT D816V mutant. Assays were generated using human recombinant KIT proteins: non-activated for the autoinhibited state, activated for the non-autoinhibited state and activated with the D816V mutation for the mutant form.

KIT phosphorylation status was evaluated by LC-MS/MS analysis, which revealed a differential phosphorylation pattern between the non-activated and activated KIT proteins (**Figures 2, S1 and S2**). The absence of phosphorylated peptides in the JM and kinase domains of non-activated KIT is consistent with its autoinhibited conformation. In contrast, activated KIT showed phosphorylation at Y547, Y553, Y568, Y570, Y703, Y721, Y823, Y900 and Y936. Interestingly, the detection of both phosphorylated and non-phosphorylated Y823 suggests that KIT proteins exist in an equilibrium between the active and inactive, non-autoinhibited states. Of note, both the autoinhibited and non-autoinhibited KIT proteins exhibited a phosphorylation pattern that mirrors those previously observed in cellular settings^4–5^, further supporting the physiological relevance of these assays.

**Figure 2:**
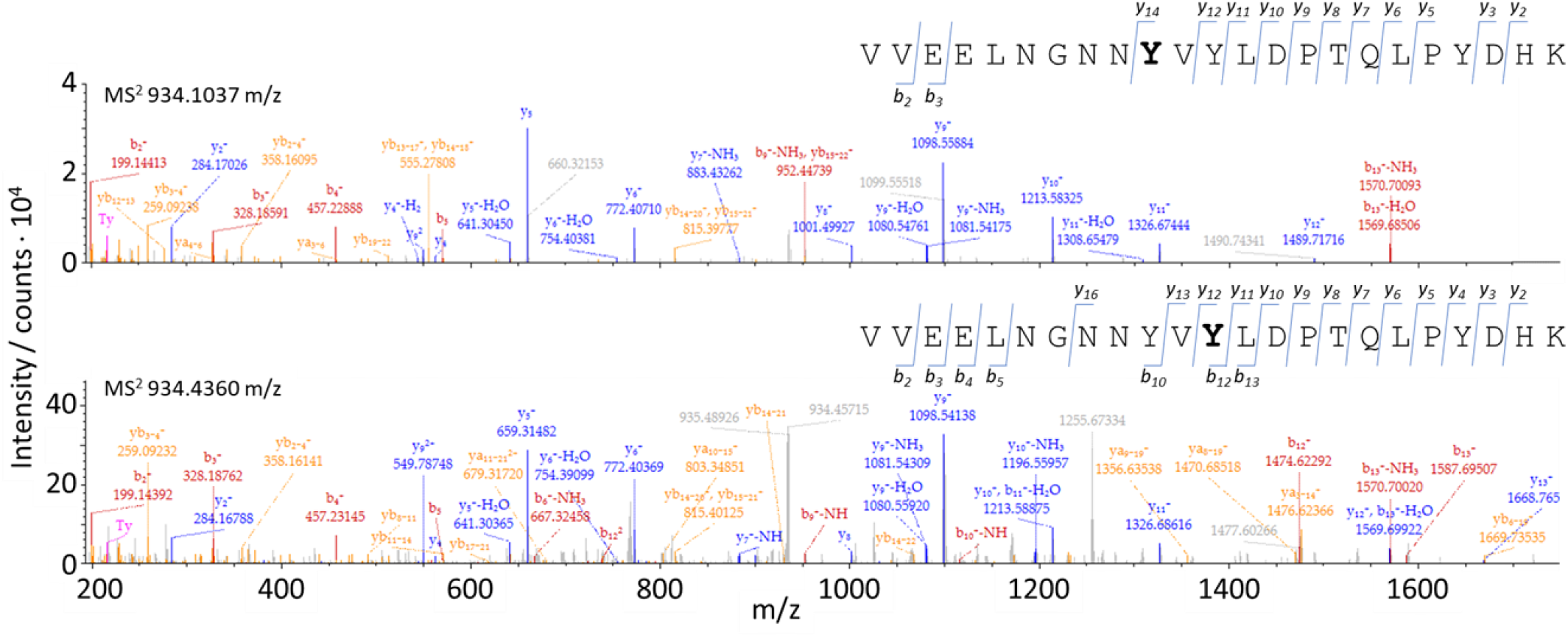
Non-autoinhibited KIT phosphorylated peptides identified by LC-MS/MS. MS^2^ fragmentation spectrum from precursor ions 934.1037 m/z (Top) and 934.4360 (Bottom) showing fragment ion ascription to the main fragmentation series: y, derived from the peptide C-terminus (blue); and b, derived from the peptide N-terminus (red). Internal fragment, immonium and precursor ions are indicated in orange, pink and green, respectively. Phosphorylated tyrosine residues pY568 (Top) and pY570 (Bottom) from the JM domain are highlighted in bold.

A high-throughput kinetic screen was subsequently performed on a library of 172 kinase inhibitors (**Table S1**) by testing their interactions with the different activation states of the KIT kinase domain. This library included 70 FDA- and NMPA-approved drugs (**Table S2**), 67 clinical candidates and 35 tool compounds currently in the discovery phase. In total, we measured 516 drug-target interactions and generated 799 individual parameters (291 affinity values and 508 kinetic values, **supplementary spreadsheet-Kinetic Profiling**). A good agreement (r = 0.86 for *K*_*d*_ and *k*_*on*_, and r = 0.84 for *k*_*off*_) was found between our experimental data and literature values (**Figure S3**).^1,9, 30–32^

### Conformational Plasticity of KIT as a Determinant of Inhibitor Binding Kinetics

The kinetic constants measured in this study exhibited a remarkably wide range, spanning from 4 to 2.6×10^7^ M^−1^ s ^−1^ for *k*_*on*_ and from 3.3×10^−6^ to 4.8×10^−1^ s^−1^ for *k*_*off*_. Residence times varied accordingly, from just 2 seconds to over 3.5 days. Binding affinities also spanned from 3.8×10^−11^ to 1.8×10^−5^ M.

Our dataset uncovered clear kinetic patterns linked to different KIT conformations, offering useful insights for improving drug design. Generally, inhibitors targeting KIT D816V showed a stronger correlation between affinity and *k*_*on*_ (r = −0.75, p < 0.0001) than with k_off_ (r = 0.31, p = 0.003), consistent with earlier findings (**Figure 3**).^16, 30^

**Figure 3.**
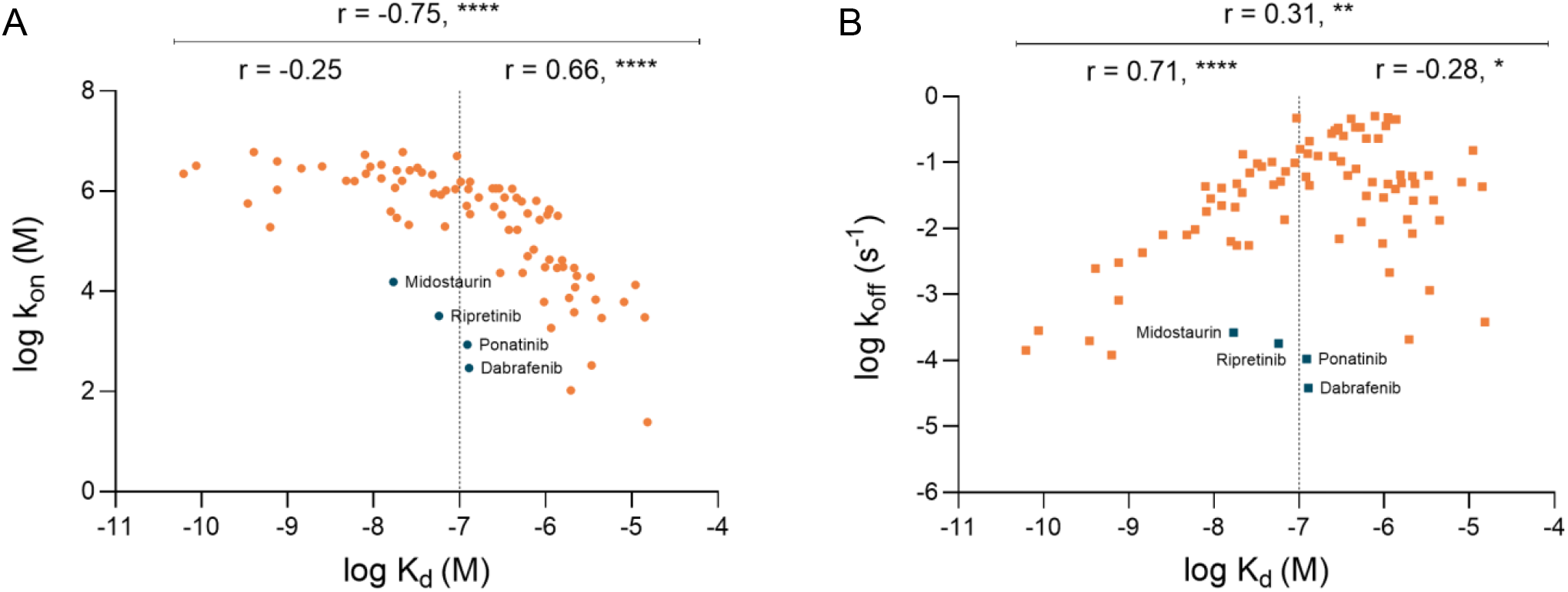
Comparison of association (A) and dissociation (B) rate constants with affinity values measured for KIT D816V. Only parameters within the assay’s detection limits are shown. Spearman correlation coefficients (r) and p-values for the entire dataset and two affinity groups (indicated by the lines) are provided above the graphs. *, p < 0.05; **, p < 0.01; ***, p < 0.001; ****, p < 0.0001.

However, this relationship was reversed for inhibitors with affinities below 100 nM, where off-rates became the key determinant of affinity (r = 0.71, p < 0.0001) while on-rates leveled off (r = −0.25, p = 0.15) near the diffusion limit (1×10^6^-1×10^7^ M^−1^s^−1^). This indicates that in the case of advanced inhibitors affinity is enhanced primarily by reducing *k*_*off*_, leading to longer residence time on KIT D816V. Interestingly, four FDA-approved inhibitors—dabrafenib, ponatinib, ripretinib and midostaurin—deviated from the general trend. All exhibited significantly extended residence times (64-434 min), suggesting that sustained target engagement may play a greater role in their efficacy than affinity alone. This distinctive behavior may reflect their ability to stabilize inactive conformations of KIT D816V. Consistent with this notion, the X-ray crystal structure of B-RAF bound to dabrafenib (PDB: 5CSW) shows the inhibitor engaging a inactive (DFG-in, αC-helix out) conformation,^33^ while ripretinib preferentially stabilizes the DFG-out conformation of the active, non-autoinhibited KIT.^32^

A strikingly different pattern emerged for inhibitors targeting the non-autoinhibited state of KIT (**Figure S4**), characterized by weaker correlation with *k*_*on*_ (r = −0.36, p = 0.0003) and stronger correlation with *k*_*off*_ (r = 0.61, p < 0.001). The increased variability in this dataset likely reflects the coexistence of active and inactive forms within the non-autoinhibited state. Restricting the analysis to inhibitors annotated in the KLIFS database^34^ as targeting the DFG-in (αC-helix in or αC-helix out) conformation, revealed a correlation pattern that closely resembled that of KIT D816V (**Figure S4**,**B**): affinity was driven by *k*_*on*_ for moderate binders (r = −0.67, p < 0.001) and by *k*_*off*_ for the most potent inhibitors (r = 0.37, p = 0.01). Conversely, inhibitors that stabilize DFG-out states **(Figure S4**,**C)**—regardless of αC-helix position—followed a distinct kinetic pattern, with affinity predominantly determined by *k*_*on*_ (r = −0.88, p < 0.0001) and showing no measurable contribution from *k*_*off*_ (r = −0.1, p = 0.65).These observations suggest divergent optimization strategies depending on the target conformation: while enhancing residence time is key for inhibiting the active conformation of non-autoinhibited KIT and KIT D816V, accelerating association rates may be more effective against the inactive non-autoinhibited form.

Building on these general trends, we next focused on high-affinity inhibitors across KIT activation states to examine how conformational differences influence kinetic behavior (**Figure 4A**). In the autoinhibited conformation, only 9% of inhibitors bound with high affinity (minimum *K*_*d*_: 2 nM). This limited binding is consistent with the structural constraints imposed by the JM-in configuration, which occludes the ATP-binding site and restricts the G-loop and αC-helix from moving into their active positions. Supporting this, association rates were the lowest across all conformational states (median *k*_*on*_: 8.7×10^4^ M^−1^s^−1^). Once bound, inhibitors formed stable complexes, with a median residence time of 17 minutes (95% CI: 1.4–139 min), a kinetic signature characteristic of a buried binding pocket.

**Figure 4:**
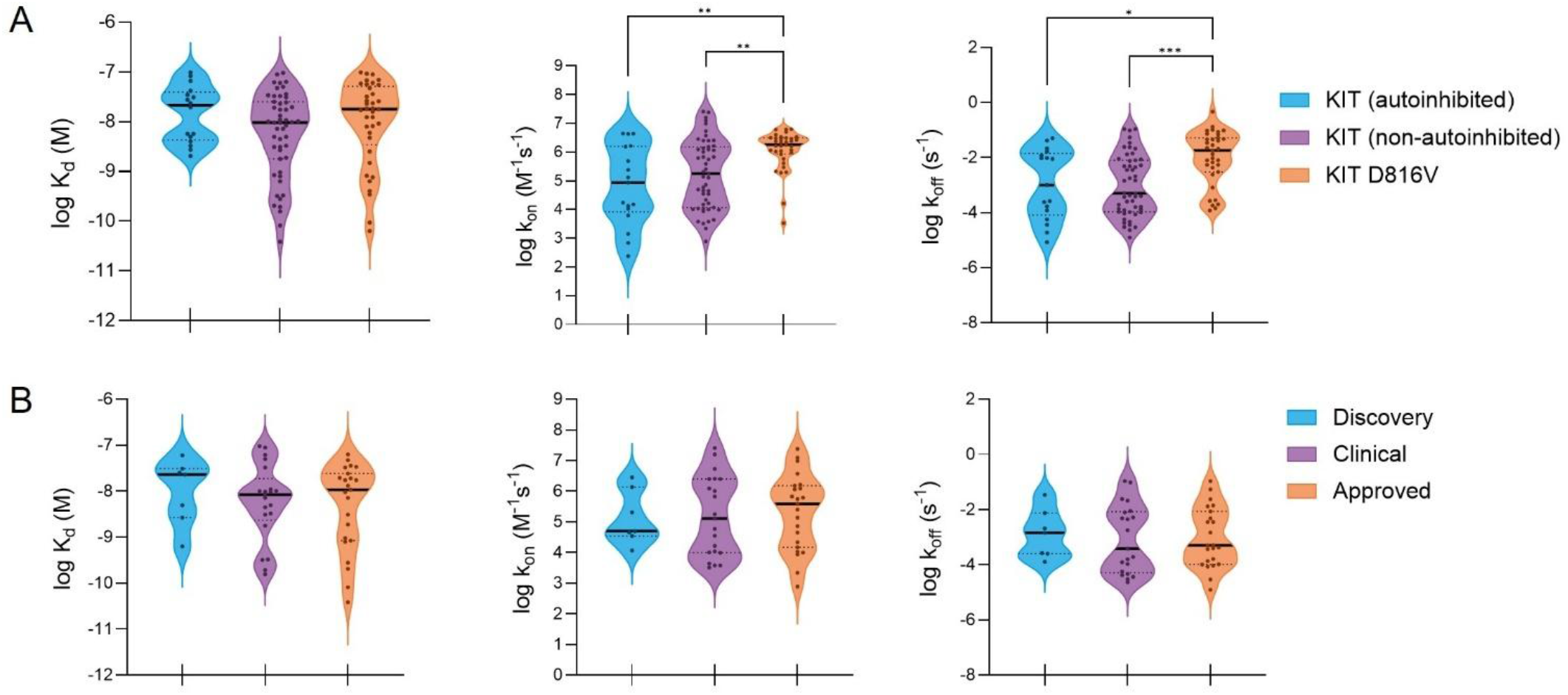
Kinetic and affinity parameters across KIT activation states. (A) Violin plots illustrate log-values of *K*_*d*_, *k*_*on*_ and *k*_*off*_ for inhibitors (*K*_*d*_ ≤ 100 nM) that interact with one or more activation states. (B) Data grouped by clinical development phase for inhibitors targeting the non-autoinhibited state. Only inhibitors within detection limits are shown. *, p < 0.05; **, p < 0.01; ***, p < 0.001.

Transition to the non-autoinhibited state, involving major rearrangements of the JM domain and A-loop along with coordinated movements of the G-loop and αC-helix, enables KIT to adopt both active and inactive conformations. Correspondingly, inhibitor engagement improved markedly: 30% of compounds bound with high affinity (0.039 nM) and faster association rates (median kon: 1.8×10^5^ M^−1^s^−1^). Interestingly, the dissociation rate remained comparable to that of the autoinhibited state (median *k*_*off*_: 5.0×10^−4^ s^−1^), yielding similar residence times (median: 33 min; 95% CI: 4.7–85 min).

The D816V mutation introduces significant conformational changes that increase flexibility within the ATP-binding pocket and enhances coordinated motions between the G-loop and the αC-helix. While only 22% of inhibitors achieved high-affinity binding (minimum *K*_*d*_: 0.063 nM), the fastest on-rate (median *k*_*on*_: 1.8×10^6^ M^−1^s^−1^) was observed. However, this increased flexibility came at the expense of stability, resulting in the highest off-rate (median *k*_*off*_: 1.8×10^−2^ s^−1^) and the shortest residence time (median: 0.9 min; 95% CI: 0.4–3 min).

### Clinically Effective Drugs Display Long-lasting Inhibition with KIT

To investigate how kinetic parameters evolve throughout drug development, we analyzed inhibitors targeting the non-autoinhibited state of KIT, the conformation with the most comprehensive dataset across all development stages (**Figure 4B**). Our analysis revealed a trend of progressively tighter binding as drugs progress through development stages, driven by a substantial rise in median *k*_*on*_ (from 5.0×10^4^ M^−1^s^−1^ in tool compounds to 3.9×10^5^ M^−1^s^−1^ in approved drugs). Consistent with previous studies,^30,35^ clinical candidates and approved drugs demonstrated considerably longer residence times (43 and 33 min, respectively) compared to the shorter interactions (9.7 min) seen with compounds still in the discovery phase. This shift underscores the rising importance of residence time, pointing to kinetic optimization as a key component in developing effective therapies. It should be noted, however, that clinical approval is determined by a complex interplay of factors—including safety, pharmacokinetics, and overall pharmacodynamic behavior—and that not all approved drugs necessarily maximize on target affinity and/or kinetics.

This trend became even more evident when examining the kinetic behavior of approved therapies for KIT-driven diseases across activation states (**Figure 5B**). With the exception of sunitinib, all exhibited prolonged residence times, highlighting the therapeutic value of sustained target engagement. In SM, midostaurin and avapritinib, remained bound to KIT D816V for over an hour. In GIST, imatinib—the frontline therapy—stood out with a 182-minute residence time on non-autoinhibited KIT. This pattern extended to ripretinib, which exhibited exceptionally long residence times—28 h on non-autoinhibited KIT and 91 min on KIT D816V—a kinetic profile that underpinned its efficacy as a fourth-line therapy. Cabozantinib, with a 107-minute residence time, showed promise in both preclinical and clinical GIST studies. Even pazopanib, despite its broader target spectrum and 43-minute residence time, demonstrated efficacy in imatinib- and sunitinib-resistant GISTs.^4,11^

**Figure 5:**
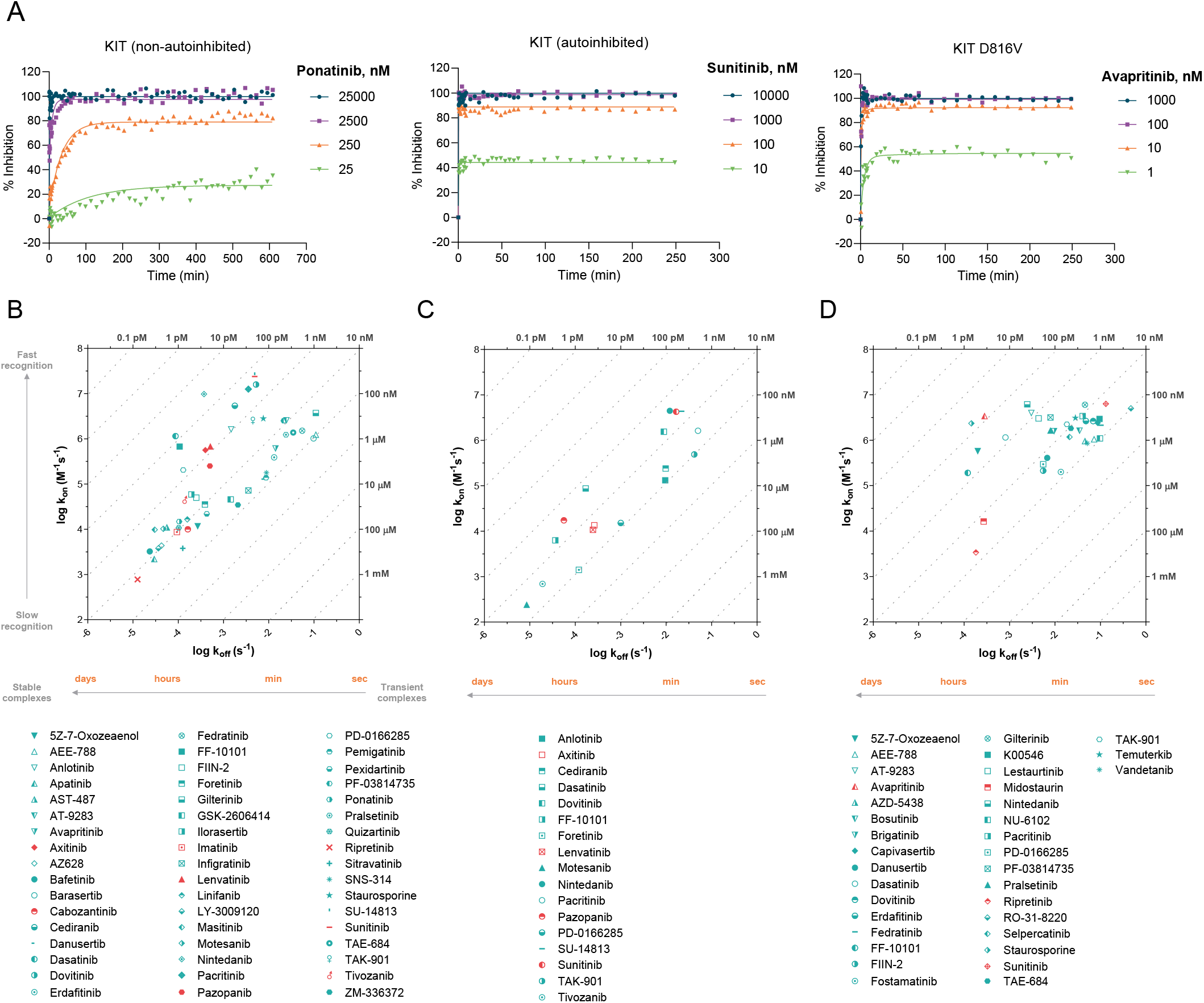
Binding kinetics of KIT-targeting inhibitors. (A) Representative kinetic curves illustrating distinct kinetic profiles: ponatinib (non-autoinhibited), sunitinib (autoinhibited) and avapritinib (KIT D816V). (B–D) Log-scale on/off-rate plots for inhibitors with *K*_*d*_ ≤ 100 nM against (B) non-autoinhibited, (C) autoinhibited and (D) D816V KIT. Dashed lines indicate *K*_*d*_ values. Drugs approved for diseases where KIT is a critical driver are shown in red.

The clinical efficacy of KIT inhibitors with prolonged residence times may be attributed to their ability to maintain inhibition throughout the receptor’s lifespan. Inhibitors that stay bound long enough to outlast receptor turnover help prevent transient signaling that could otherwise promote oncogenic activity. In NIH 3T3 cells, SCF-stimulated non-autoinhibited KIT internalizes and degrades within 30-60 min, while autoinhibited KIT levels remain stable for up to 4 h without ligand stimulation. The D816V mutant, despite lacking stimulation, degrades within 1-2 h.^36^

### Kinase Activation Features and Binding Site Location Control Binding Kinetics

Inhibitor binding modes are shaped by both the structural differences between active and inactive kinase forms and the specific interactions between the inhibitor and the kinase. The KLIFS database^34^ maps these interactions by classifying inhibitors according to their engagement with specific structural pockets (front cleft, gate area and back cleft) and subpockets. Type I inhibitors bind to the ATP-binding site in the active conformation, with the DFG motif and αC-helix facing inward (DFG-in, αC-helix in), primarily occupying the front cleft. In contrast, type II inhibitors target the inactive DFG-out conformation, where the DFG motif points away from the active site pushing the αC-helix outwards (αC-helix out), exposing hydrophobic regions in the back cleft. Type I½ inhibitors, a subtype of type I, bind to an inactive state with the DFG-in, αC-helix out configuration, occupying the front cleft while extending into the gate and back cleft. Importantly, type I½ and II inhibitors can be divided into A and B subclasses, depending on whether they extend past the gatekeeper residue into the back cleft (A) or remain confined to the front cleft (B). This structural difference is reflected in their kinetics, with type B inhibitors typically showing shorter residence times than type A compounds. Allosteric compounds bind exclusively to the back cleft without interacting with the ATP-binding pocket: type III inhibitors bind adjacent to the ATP-binding site while type IV inhibitors target remote regions outside the catalytic site. Additionally, type VI inhibitors form covalent, often irreversible, bonds with their targets. Understanding these binding modes is essential for designing effective therapies against kinases such as KIT, which can adopt multiple activation states within cells.^4,33,37.^

Analysis of 167 compounds with known binding modes^33–65^ (**Table S3**) confirmed the ability of KINETICfinder to detect a broad range of inhibitor modalities (**Figure 6**). The platform accurately characterized type I (38%), I½ (20%), II (23%), III (5%), IV (1%) and VI (14%) inhibitors, which is further supported by previous LIMK1 studies showing a strong correlation between type III inhibitor affinity and biochemical and cellular data.^66^ Such diversity reflects the pharmaceutical strategy of targeting multiple kinase conformations to enhance therapeutic efficacy.

**Figure 6:**
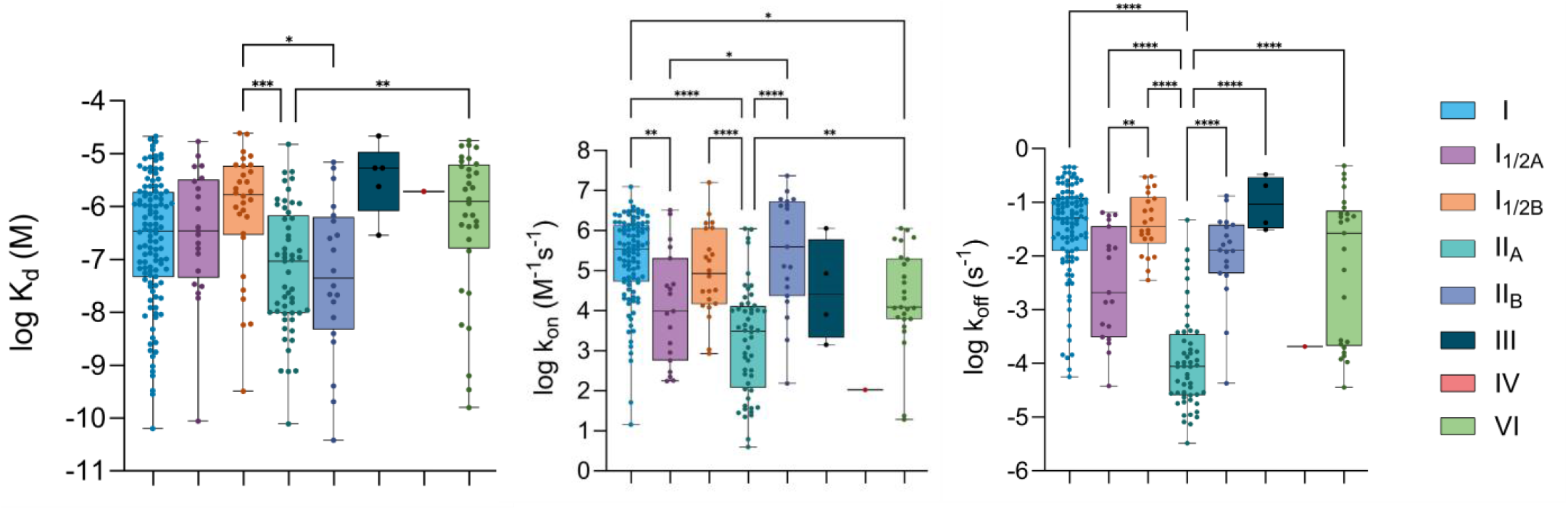
Box-and-whisker plots depicting log-distributions of *K*_*d*_, *k*_*on*_ and *k*_*off*_ for inhibitors grouped by binding modes. Only inhibitors within detection limits are shown. *, p < 0.05; **, p < 0.01; ***, p < 0.001; ****, p < 0.0001.

A closer look at residence times identified distinct kinetic signatures among inhibitor types. Type I, I½B, IIB, III and VI inhibitors primarily formed short-lived drug-target complexes with median residence times of 0.2-1 min. For type VI inhibitors, this apparent similarity to reversible classes reflects a key limitation of our dataset: none of the covalent candidates were originally designed to target KIT, and most did not show time-dependent inhibition, indicating non-covalent, off-target behavior. Even so, a small subset showed clear time-dependent inhibition, consistent with potential covalent engagement, contributing to the wide 95% confidence interval of their median residence time (0.3–66 min). In contrast, type I½A and IIA inhibitors typically established long-lasting interactions (median: 8 and 189 min, respectively). However, notable exceptions were observed. Type I inhibitors pazopanib and midostaurin, along with the type I½ inhibitor avapritinib (**Table 1**) and IIB nintedanib, displayed an unexpectedly prolonged residence time.

**Table 1.** Kinetic and affinity profiles of kinase inhibitors across KIT states.

| KIT (autoinhibited) |  |  |  |  |  |
| --- | --- | --- | --- | --- | --- |
| Compounds | Binding mode | $k_{on}$ (M<sup>-1</sup>s<sup>-1</sup>) | $\tau$ (min) | $K_d$ (nM) | N |
| Avapritinib | I<sub>1/2</sub> | (2.7±0.5)×10<sup>4</sup> | 1.8±0.1 | 349±37 | 2 |
| Axitinib | IIA | (1.3±0.1)×10<sup>4</sup> | 64±2 | 20±0.4 | 2 |
| Bosutinib | IIB | (1.5±0.3)×10<sup>2</sup> | 388±1 | 292±60 | 2 |
| Cabozantinib | IIA | (3.4±2.0)×10<sup>2</sup> | 683±260 | 101±29 | 2 |
| Dasatinib monohydrate | IIA | (8.7±1.4)×10<sup>4</sup> | 107±39 | 2.0±0.7 | 5 |
| GSK-2606414 | I<sub>1/2A</sub> | (8.0±1.4)×10<sup>2</sup> | 78±8 | 272±18 | 2 |
| Imatinib mesylate | IIA | (2.4±0.7)×10<sup>1</sup> | 624±144 | 1216±81 | 2 |
| Lenvatinib | IIA | (1.1±0.1)×10<sup>4</sup> | 72±15 | 24±0.8 | 2 |
| Linifanib | IIA | (1.0±0.2)×10<sup>2</sup> | 1338±37 | 120±6 | 2 |
| Masitinib | IIA | (3.8±0.4)×10<sup>1</sup> | 1542±71 | 288±47 | 2 |
| Midostaurin hydrate | I | NE | NE | >10000 | 2 |
| Nintedanib | IIB | (4.5±0.9)×10<sup>6</sup> | 1.4±0.3 | 2.7±0.1 | 3 |
| Pazopanib | I | (1.7±0.3)×10<sup>4</sup> | 298±38 | 3.3±0.1 | 2 |
| Pexidartinib hydrochloride | IIA | (1.5±0.1)×10<sup>2</sup> | 185±18 | 599±88 | 2 |
| Ponatinib | IIA | (6.6±0.8)×10<sup>1</sup> | 642±9 | 402±45 | 2 |
| Ripretinib | IIA | (6.2±0.9)×10<sup>0</sup> | 2054±202 | 1359±319 | 2 |
| Staurosporine | I | (8.1±0.1)×10<sup>4</sup> | 0.19±0.02 | 1106±127 | 2 |
| Sunitinib malate | IIB | (4.2±0.8)×10<sup>6</sup> | 1.0±0.1 | 4.0±0.9 | 3 |
| KIT (non-autoinhibited) |  |  |  |  |  |
| Compounds | Binding mode | $k_{on}$ (M<sup>-1</sup>s<sup>-1</sup>) | $\tau$ (min) | $K_d$ (nM) | N |
| Avapritinib | I<sub>1/2</sub> | (6.2±1.1)×10<sup>5</sup> | 1.2±0.2 | 25±8 | 2 |
| Axitinib | IIA | (5.6±2.4)×10<sup>5</sup> | 44±11 | 0.79±0.22 | 3 |
| Bosutinib | IIB | (1.7±0.3)×10<sup>4</sup> | 3.9±0.4 | 263±73 | 2 |
| Cabozantinib | IIA | (1.0±0.4)×10<sup>4</sup> | 107±24 | 18±3 | 2 |
| Dasatinib monohydrate | IIA | (1.2±0.3)×10<sup>6</sup> | 192±24 | 0.082±0.024 | 5 |
| GSK-2606414 | I<sub>1/2A</sub> | (4.6±0.5)×10<sup>4</sup> | 12±3 | 31±4 | 2 |
| Imatinib mesylate | IIA | (8.7±0.4)×10<sup>3</sup> | 182±20 | 11±2 | 2 |
| Lenvatinib | IIA | (6.7±1.9)×10<sup>5</sup> | 33±5 | 0.85±0.34 | 2 |
| Linifanib | IIA | (1.7±0.7)×10<sup>4</sup> | 113±32 | 10±3 | 4 |
| Masitinib | IIA | (9.7±0.4)×10<sup>3</sup> | 607±181 | 3.1±0.8 | 2 |
| Midostaurin hydrate | I | (1.1±0.4)×10<sup>5</sup> | 0.77±0.08 | 221±72 | 3 |
| Nintedanib | IIB | (9.8±1.7)×10<sup>6</sup> | 46±7 | 0.039±0.007 | 3 |
| Pazopanib | I | (2.5±0.7)×10<sup>5</sup> | 43±21 | 1.9±0.4 | 2 |
| Pexidartinib hydrochloride | IIA | (2.2±0.2)×10<sup>4</sup> | 41±7 | 19±2 | 2 |
| Ponatinib | IIA | (1.5±0.2)×10<sup>4</sup> | 160±7 | 7.2±0.5 | 2 |
| Ripretinib | IIA | (7.7±0.6)×10<sup>2</sup> | 1717±836 | 17±9 | 2 |
| Staurosporine | I | (2.8±0.2)×10<sup>6</sup> | 2.2±0.4 | 2.7±0.4 | 7 |
| Sunitinib malate | IIB | (2.4±0.6)×10<sup>7</sup> | 3.6±0.9 | 0.20±0.04 | 2 |

| KIT D816V |  |  |  |  |  |
| --- | --- | --- | --- | --- | --- |
| Compounds | Binding mode | $k_{on}$ ($M^{-1}s^{-1}$) | $\tau$ (min) | $K_d$ (nM) | N |
| Avapritinib | I<sub>1/2</sub> | $(3.4 \pm 0.9) \times 10^6$ | 60±7 | 0.094±0.037 | 5 |
| Axitinib | IIA | $>3.7 \times 10^5$ | <0.033 | 1376±165 | 2 |
| Bosutinib | IIB | $(1.6 \pm 0.5) \times 10^6$ | 0.49±0.07 | 23±4 | 2 |
| Cabozantinib | IIA | NE | NE | >25000 | 2 |
| Dasatinib monohydrate | IIA | $(1.2 \pm 0.3) \times 10^6$ | 21±1 | 0.81±0.34 | 4 |
| GSK-2606414 | I<sub>1/2A</sub> | NE | NE | >25000 | 2 |
| Imatinib mesylate | IIA | $(4.0 \pm 1.1) \times 10^3$ | 2.5±1.3 | 2227±685 | 2 |
| Lenvatinib | IIA | NE | NE | >2500 | 2 |
| Linifanib | IIA | NE | NE | >25000 | 2 |
| Masitinib | IIA | $(2.9 \pm 0.3) \times 10^3$ | 1.3±0.2 | 4618±355 | 2 |
| Midostaurin hydrate | I | $(1.6 \pm 0.5) \times 10^4$ | 66±15 | 17±3 | 3 |
| Nintedanib | IIB | $(6.1 \pm 0.9) \times 10^6$ | 6.9±0.7 | 0.41±0.06 | 6 |
| Pazopanib | I | $(4.4 \pm 1.6) \times 10^5$ | 0.06±0.02 | 776±46 | 2 |
| Pexidartinib hydrochloride | IIA | NE | NE | >2500 | 2 |
| Ponatinib | IIA | $(9.1 \pm 2.4) \times 10^2$ | 159±11 | 122±24 | 2 |
| Ripretinib | IIA | $(3.4 \pm 0.9) \times 10^3$ | 91±2 | 59±15 | 2 |
| Staurosporine | I | $(2.4 \pm 0.7) \times 10^6$ | 123±31 | 0.063±0.012 | 6 |
| Sunitinib malate | IIB | $(6.3 \pm 2.0) \times 10^6$ | 0.13±0.01 | 23±9 | 2 |

When analyzing *k*_*on*_, an inverse relationship to the dissociation rate constant was observed. Type I and IIB inhibitors exhibited significantly faster binding kinetics, with a median *k*_*on*_ of 3.4×10^5^ and 4.0×10^5^ M^−1^s^−1^, compared to the slower *k*_*on*_ of 3.1×10^3^ and 9.8 x10^3^ M^−1^s^−1^ observed for type IIA and I½A inhibitors, respectively. These findings suggest that while type I and IIB inhibitors rapidly interact with their targets, type I½A and IIA inhibitors rely on slower association rates and longer residence times. Notably, both type IIA and IIB inhibitors demonstrated the highest binding affinities for KIT among all inhibitor classes tested.

Further analysis based on key structural activation features highlighted additional kinetic fingerprints (**Figure 7A and 7B**). Inhibitors that stabilize the DFG-out conformation—whether in the αC-helix in or αC-helix out orientation— exhibited higher affinities and residence times extending into the hours range. In contrast, inhibitors targeting the DFG-in, αC-helix in and DGF-out-like, αC-helix in conformations shared similar kinetic profiles, rapid on-rates and short residence times on a second time scale. Binding site selection further influenced kinetic trends. In general, front-cleft inhibitors exhibited fast on- and off-rates whereas back-cleft inhibitors associated more slowly and displayed prolonged residence times, probably due to deeper integration into the kinase structure.

**Figure 7:**
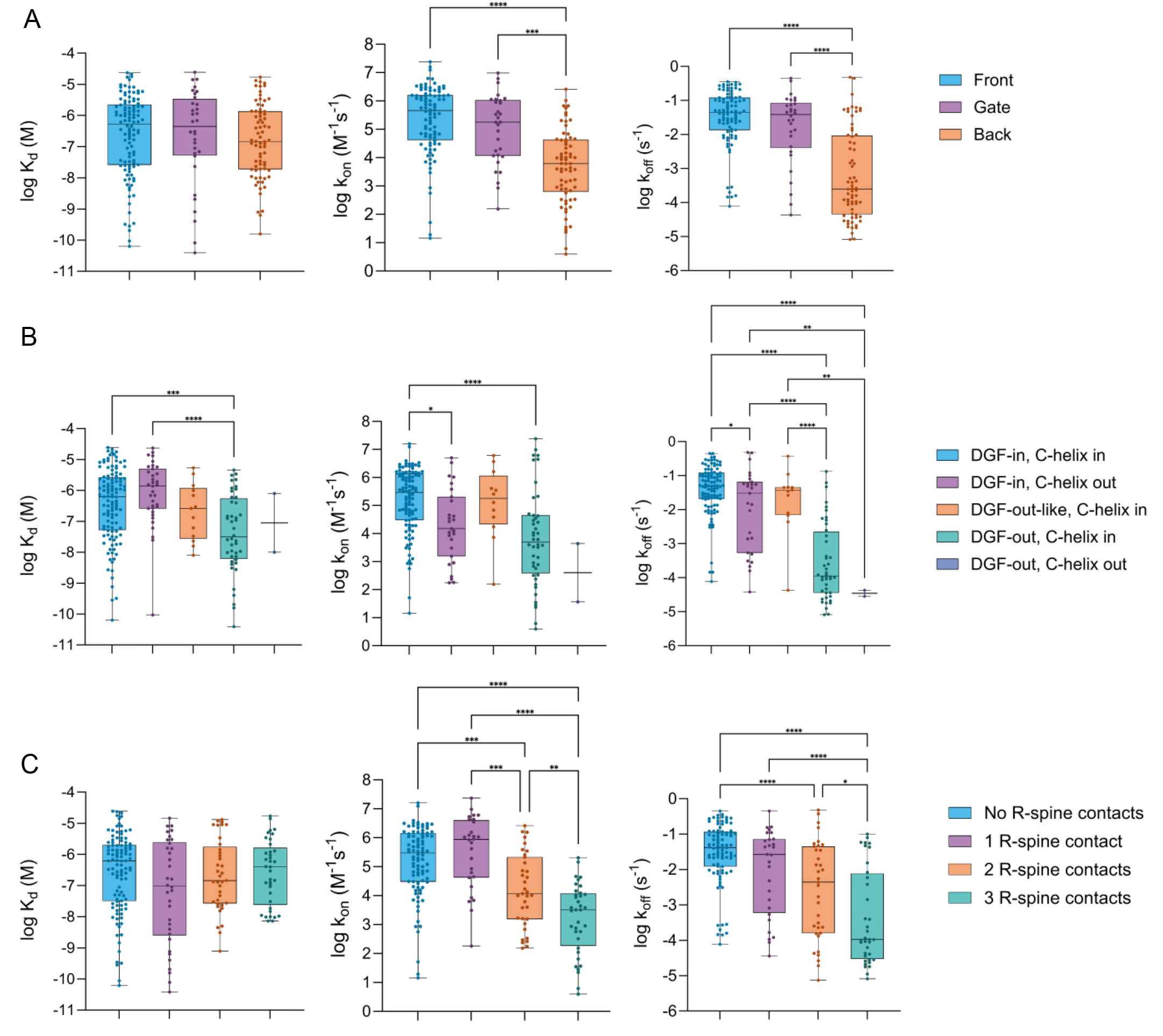
Structural determinants of inhibitor binding kinetics. Box-and-whisker plots depicting log-distributions of *K*_*d*_, *k*_*on*_ and *k*_*off*_ for inhibitors grouped by (A) KLIFS-defined pockets, (B) DFG motif orientation and αC-helix position and (C) R-spine engagement (reflecting increasing contact from none to one, two, or three R-spine residues). Only inhibitors within detection limits are shown. *, p < 0.05; **, p < 0.01; ***, p < 0.001; ****, p < 0.0001.

Another key dynamic element is the regulatory spine (R-spine). This four-residue assembly (RS1–RS4) stabilizes the DFG motif and αC-helix via hydrophobic interactions, with proper R-spine alignment being necessary for catalytic activity. Previous studies have demonstrated that residence times of type I½ inhibitors of p38α MAPK^67^ and type II inhibitors of BTK can be modulated by altering R-spine interactions.^68^ To investigate whether this regulatory mechanism extends to KIT, we examined inhibitor interactions with the R-spine using the KLIFS database (**Figure 7C**). Our analysis revealed a clear pattern linking R-spine engagement to kinetic behavior. Inhibitors lacking R-spine contacts displayed short residence times (median: 0.4 min), whereas those interacting with up to three R-spine residues showed markedly prolonged target engagement (median: 159 min), a nearly 400-fold difference. This extended residence time was accompanied by substantially slower association rates (median: 2.9×10^5^ for no R-spine contacts vs 3.2×10^3^ M^−1^s^−1^ for three contacts). Extending the analysis across KIT conformations (**Figure S5**) showed that R-spine interactions shape both the binding kinetics and affinity of inhibitors targeting autoinhibited and non-autoinhibited KIT. By contrast, inhibitors engaging the R-spine in KIT D816V exhibit reduced affinity due to a slower association rate.

### Faster Dissociation Rates in D816V utation Contribute to Drug Resistance

Drug resistance is often associated with a reduction in residence time, where mutations weaken drug binding by increasing the off-rate rather than affecting the on-rate. In chronic myeloid leukemia, analysis of 94 resistance-associated mutants of the ABL kinase showed that only 32% caused a ≥4-fold drop in imatinib affinity, with most displaying unchanged *K*_*d*_ but faster dissociation. Structural analysis of the N368S mutant revealed enhanced dynamics in the A-loop and αC-helix, promoting imatinib dissociation. A similar trend has been observed in resistance mutations of HIV-1 protease and reverse transcriptase, where clinical inhibitors such as amprenavir, ritonavir or saquinavir primarily lost efficacy due to decreased residence times, rather than changes in *k*_*on*_.^16, 17, 69^

Our findings (**Figure 8A, Table 1**) reinforce the role of binding kinetics in drug resistance associated with the KIT D816V mutation. We found that 43% of the inhibitors tested exhibited a significant (>3-fold) drop in affinity for KIT D816V compared to the non-autoinhibited KIT, with some showing up to a 1500-fold affinity decrease. As expected, structural analysis based on available literature data for KIT inhibitors ^9, 33, 38, 63, 64^ (**Figure 8B, Table S3**) showed that those with the greatest loss of affinity are type II compounds. These inhibitors target the inactive DFG-out conformation, characterized by the repositioning of F811 into the ATP-binding pocket to form the back cleft. This site is engaged by pexidartinib, lenvatinib, axitinib, imatinib, cabozantinib and ponatinib, which establish key contacts with F811 and E640 located at the αC-helix. However, the D816V mutation destabilizes the A-loop and αC-helix, likely disrupting these interactions and reducing drug affinity. In contrast, sunitinib and dasatinib, which bind the front cleft and gate area respectively, retained high affinity for KIT D816V although reduced relative to non-autoinhibited KIT, presumably by interacting with F811 but not E640 (**Figure 9A**). This behavior is consistent with prior *in silico* modeling of intermediate conformational states of KIT WT and KIT D816V, which suggests that sunitinib progressively loses stabilizing interactions as the activation loop shifts and becomes increasingly constrained by mutation-induced rearrangements of key structural elements.^8^

**Figure 8:**
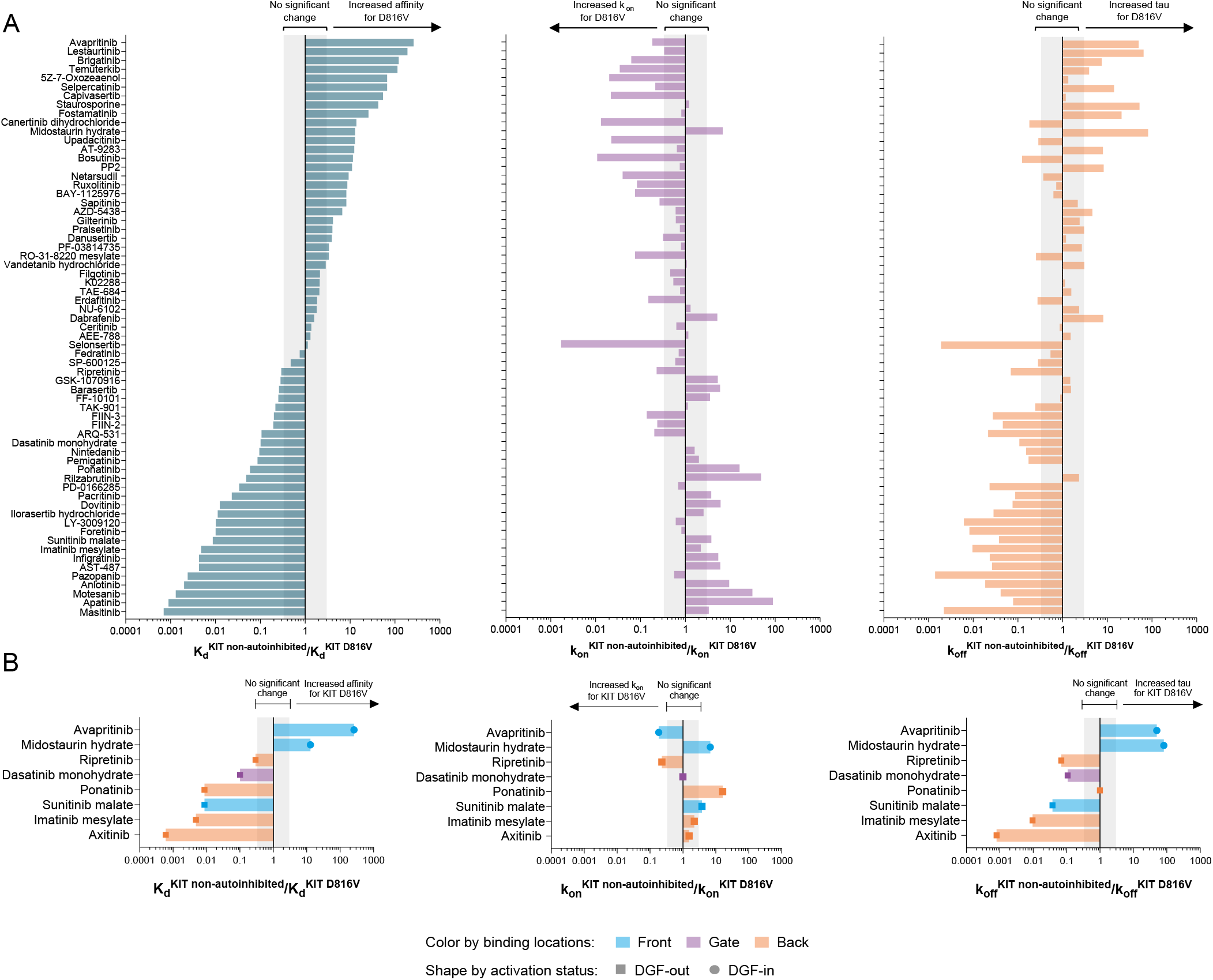
Comparative analysis of kinetic parameters between KIT non-autoinhibited and KIT D816V mutant. (A) Bar plots display fold changes in *K*_*d*_, *k*_*on*_ and *k*_*off*_ relative to KIT non-autoinhibited. (B) Analysis restricted to inhibitors with structural data for KIT, grouped by binding pocket (color) and DFG status (shape). Gray area indicates no significant change.

**Figure 9:**
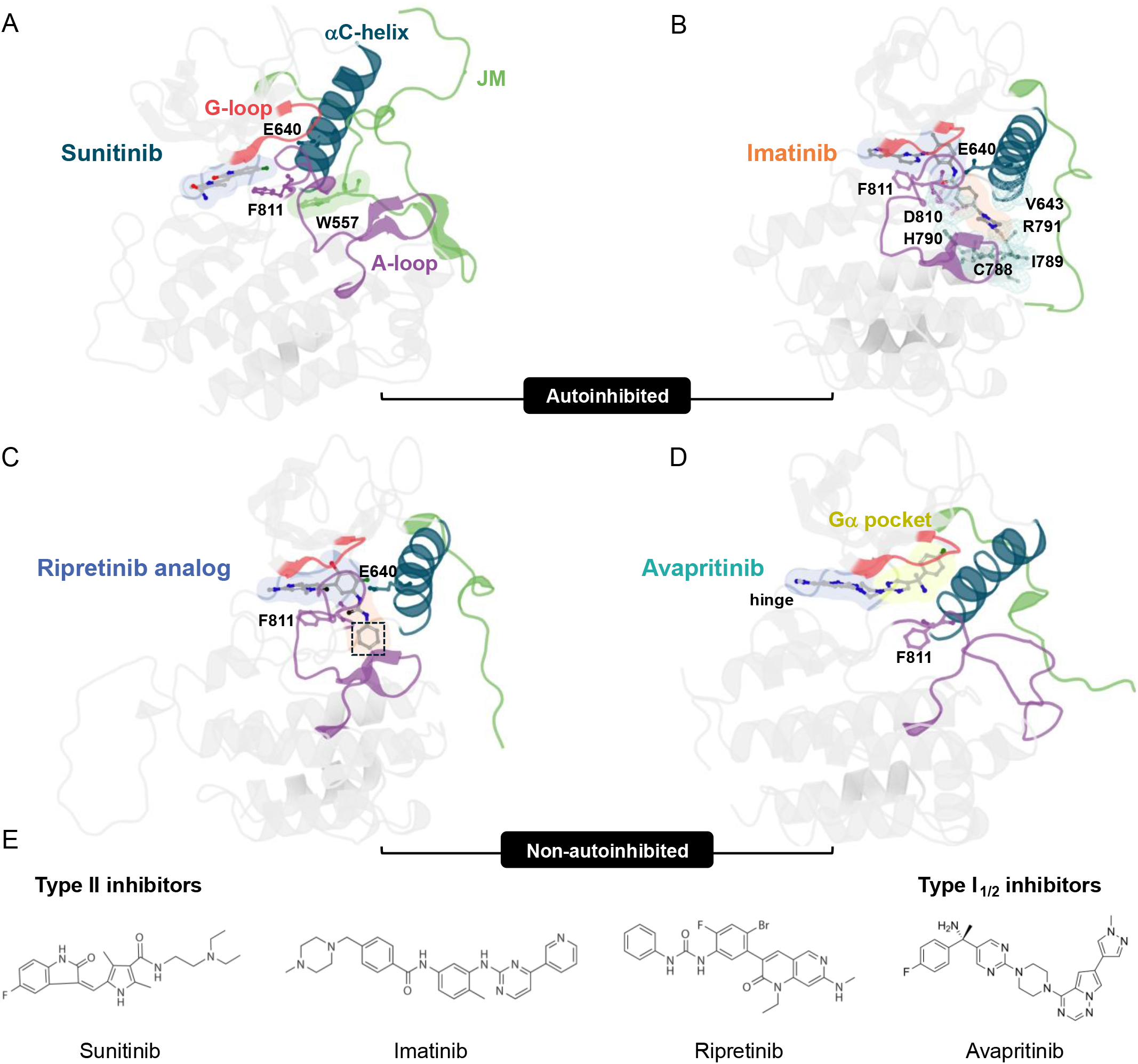
Crystal structures of type I_1/2_ and type II inhibitors bound to KIT. (A) Sunitinib binds the front cleft (violet) of autoinhibited KIT, with W557 from the JM domain (green) displacing F811 to stabilize the A-loop (purple) in a DFG-out conformation (PDB: 3G0E). (B) Imatinib occupies both front and back (orange) clefts of autoinhibited KIT (PDB: 1T46). (C) A close ripretinib analog binds non-autoinhibited KIT, with its terminal phenyl ring (black box) occupying the position normally held by W557 in the autoinhibited state, thereby stabilizing the DFG-out conformation (PDB: 6MOB). (D) Avapritinib engages non-autoinhibited KIT in a DFG-in conformation, with its fluorobenzene moiety occupying the Gα-pocket (yellow) (PDB: 8PQH). (E) Chemical structures of the inhibitors shown in panels A–D.

**Figure 10:**
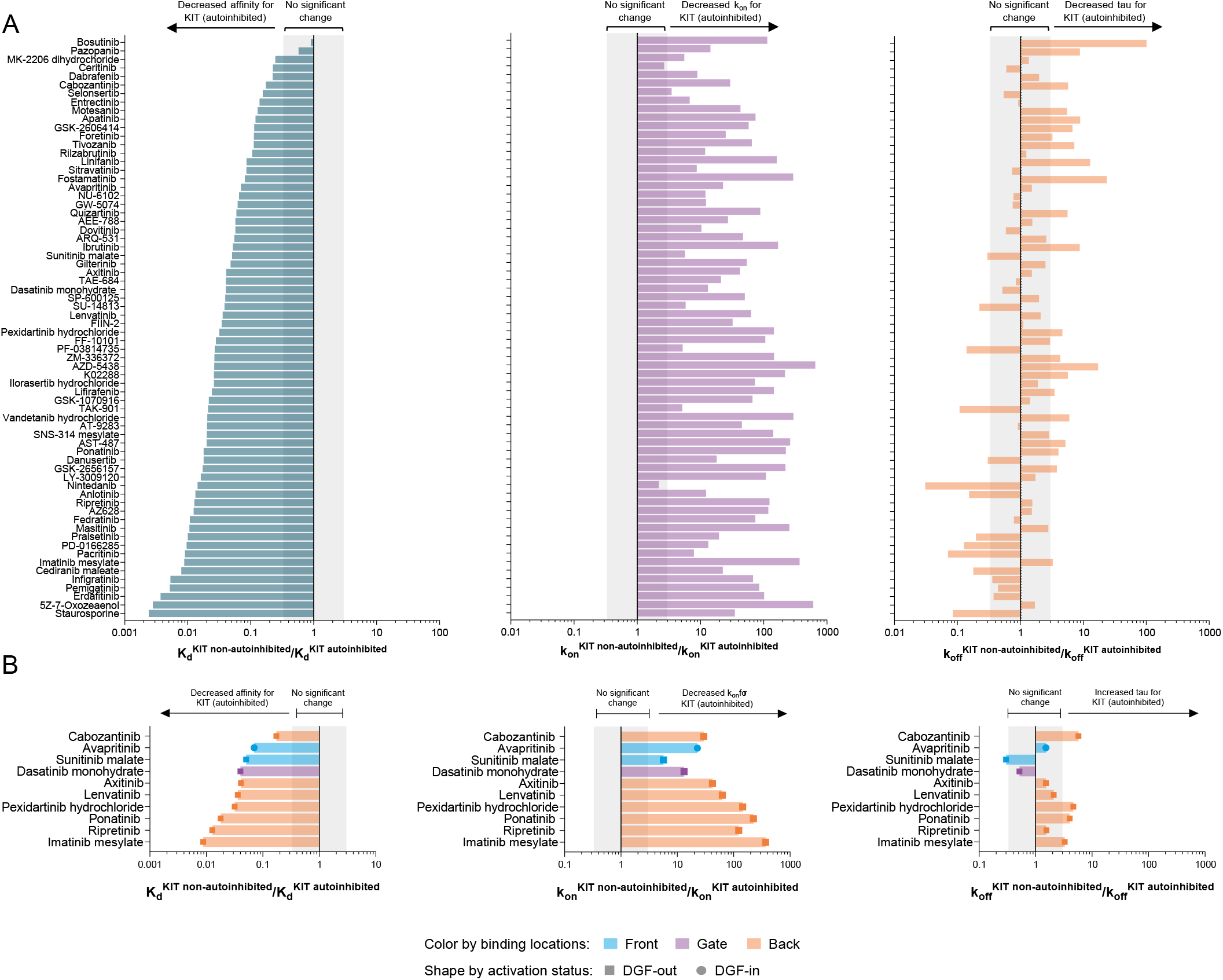
Comparative analysis of kinetic parameters between non-autoinhibited and autoinhibited KIT. (A) Bar plots show fold changes in *K*_*d*_, *k*_*on*_ and *k*_*off*_ relative to non-autoinhibited KIT. (B) Analysis restricted to inhibitors with structural data for KIT, grouped by binding pocket (color) and DFG status (shape). Gray area indicates no significant change.

In 96% of the cases of inhibitors tested, this loss in binding affinity was primarily driven by a substantial increase in *k*_*off*_, reaching more than 700-fold. Structural data (**Table S3**) further confirmed that inhibitors binding the DFG-out conformation, regardless of their binding site, experience the most pronounced reductions in residence time. This kinetic resistance was evident in kinase inhibitors such as masitinib, pazopanib, imatinib, dasatinib and sunitinib (**Table 1**), reinforcing clinical observations that SM patients harboring KIT D816V often exhibit primary resistance to imatinib and masitinib. A similar trend was observed with the ABL1 inhibitor dasatinib, although it retained high affinity for the mutant its clinical efficacy in a phase II trial remained limited.^4,7^ A plausible explanation is that its residence time for KIT D816V (~21 min) is insufficient to outlast receptor turnover, resulting in transient inhibition that fails to produce a durable therapeutic effect. In contrast, ponatinib, which had a marked drop in affinity, remained effective in KIT D816V-positive mast cells and xenograft models^70^, which may be attributed to its prolonged residence time when binding to the D816V mutant (159 min). These findings underscore the importance of optimizing residence time in the design of next-generation inhibitors targeting KIT D816V to overcome resistance mechanisms driven by increased *k*_*off*_ rates.

### Prolonged Residence Times in KIT D816V Enhance Drug Efficacy

In contrast to the predominant loss of affinity observed in most inhibitors, 35% displayed enhanced binding with KIT D816V, suggesting that the mutant conformation can favorably accommodate specific compounds. Structural data for KIT-targeting inhibitors (**Figure 8B, Table S3**) confirmed that the most potent inhibitors bind to the active DFG-in conformation, with affinity gains primarily driven by faster association rates—57% displayed up to a 93-fold acceleration in *k*_*on*_. Bosutinib, capivasertib and temuterkib exemplify this trend. Additionally, 35% of inhibitors showed simultaneous increases in both k_on_ and residence time, as exemplified by avapritinib, which achieved a remarkable 260-fold affinity gain driven by a 50-fold prolonged residence time and a modest 5-fold increase in *k*_*on*_. Strikingly, midostaurin uniquely compensated for a slower *k*_*on*_ with an 82-fold increase in residence time, highlighting how extended target engagement can sustain efficacy despite weaker initial binding (**Table 1**).

Previously published structural data^64^ show that avapritinib binds to the front cleft of non-autoinhibited KIT. Specifically, avapritinib interacts with the hinge region and the A-loop, while the fluorobenzene moiety of the inhibitor fits into a hydrophobic pocket in the G-loop (Gα-pocket). This pocket overlaps with the front pocket II (FP-II) region, located between the G-loop and the β3-strand, and extends toward the αC-helix. Structural evidence suggests this pocket forms only upon ligand binding, stabilizing a DFG-in, αC-helix out conformation (**Figure 9D**). A similar binding mechanism has been observed for staurosporine, where the glycosidic group engages with the G-loop in LCK, CDK2 and PKA, triggering a significant displacement of the G-loop upon staurosporine binding.^71^ Molecular docking studies further suggested that midostaurin, a staurosporine analog, binds within the front cleft including FP-II pocket of active KIT WT, forming hydrogen bonds with residues of the hinge region and G-loop through its benzoyl-substituted glycosidic group. However, unlike avapritinib, it does not interact with the A-loop.^38^

Molecular dynamics (MD) simulations of KIT D816V revealed a unique positioning of the αC-helix and folding of the G-loop, structural changes not observed in crystal structures of either the active non-autoinhibited or inactive autoinhibited KIT.^2^ In line with these predictions, superimposition of representative structures of KIT drug complexes (**Figure S6**) showed the plasticity of the KIT kinase domain.^72^ Based on our findings, we hypothesize that the D816V-associated rearrangement of the G-loop and αC-helix favors the access and accommodation of inhibitors that engage the FP-II region or extend into the adjacent Gα-pocket (e.g., avapritinib, midostaurin, staurosporine), thereby stabilizing the bound complex and producing longer residence times and higher affinity. Similar kinetic signatures identified for additional FP-II binders from KLIFS database, such as dabrafenib, pralsetinib, selpercatinib and temuterkib, pointed to a shared binding mechanism.

### Reduced association rates reveal structural barriers to inhibitor binding in autoinhibited KIT

The shift from the autoinhibited to the active, non-autoinhibited conformation of KIT requires large-scale structural movements, particularly in the JM domain, A-loop, αC-helix, and, to a lesser extent in the G-loop. In the autoinhibited state, the JM domain residues Y547 to G565 are buried in the kinase domain, with W557 displacing F811 to stabilize the A-loop in a DFG-out conformation. V559 and V560 stabilize this state by forming a hydrophobic patch with V643 and T646 on the αC-helix, and C788 and I789 on the catalytic loop.^63^ Our data show that these rearrangements substantially impair inhibitor binding (**Figure 1**). For 97% of the inhibitors tested, affinity decreased 4 to 412-fold compared to the non-autoinhibited state, mainly due to a significant reduction in the *k*_*on*_ (up to 653 times). In contrast, *k*_*off*_ remained unchanged for 51% of inhibitors, while 31% showed slower dissociation (up to 101-fold) and 18% exhibited faster *k*_*off*_ values (up to 32-fold).

To better interpret these differences in binding kinetics, we focused on sunitinib, a type IIB inhibitor with extensive structural data.^63,9^ Sunitinib binds KIT in a conformation resembling the autoinhibited apo state, slightly reorienting F811 to fit into the ATP-binding pocket. This binding mode is consistent with its interaction with autoinhibited VEGFR2 (PDB 4AGD).^73^ In both cases, the inhibitor occupies the front cleft without making direct contact with the JM domain, suggesting that sunitinib binds without displacing the JM domain from its autoinhibitory position (**Figure 9A**). Our kinetic data corroborated these structural observations: sunitinib exhibited fast association rates with both non-autoinhibited (2.4×10^7^ M^−1^s^−1^) and autoinhibited KIT (4.2×10^6^ M^−1^s^−1^), supporting a binding mode compatible with the JM-in conformation (**Table 1**). However, the autoinhibited form showed a sixfold reduction in *k*_*on*_, possibly due to DFG motif rearrangements upon inhibitor binding. This resulted in a faster *k*_*off*_ and overall lower affinity compared to the non-autoinhibited state. Other inhibitors targeting the front cleft or the gate area—such as nintedanib, pacritinib, anlotinib and cediranib—exhibited similar kinetic fingerprints, suggesting a comparable binding mechanism. In particular, cediranib, which exhibits a *k*_*on*_ of 5.3×10^6^ and 2.4×10^5^ M^−1^s^−1^ for non-autoinhibited and autoinhibited KIT, respectively, has been previously reported to be compatible with the JM-in conformation, further supporting this interpretation.^73^

Although the type IIA inhibitor imatinib also targets the autoinhibited conformation, it binds deeply within the back cleft, displacing the JM domain through direct competition. Crystallographic data^63^ demonstrates that its phenyl ring interacts with D810 and E640, while its piperazinyl ring fits into a pocket formed by V643, C788, I789, H790, R791 and D810. These interactions are incompatible with the JM-in position of autoinhibited KIT (**Figure 9B**). This structural incompatibility was reflected in imatinib’s extremely slow association rate (24 M^−1^s^−1^) and long residence time (10 h) (**Table 1**). This may reflect the significant structural rearrangement required for KIT to adopt a JM-out conformation. In contrast, binding to the non-autoinhibited state showed a 363-fold faster on-rate (8.7×10^3^ M^−1^s^−1^) and 3 times shorter residence time (3 h), indicating reduced steric hindrance from the JM domain. Similar kinetic behavior was observed in other back cleft inhibitors like ripretinib, ponatinib, pexidartinib, cabozantinib, GSK-2606414, tivozanib, linifanib and masitinib, suggesting a common binding mechanism. Linifanib and tivozanib (PDB 4ASE), which also bind very slowly to the autoinhibited state (100 and 690 M^−1^s^−1^, respectively) but markedly faster to the non-autoinhibited state (1.7×10^4^ and 4.5×10^4^ M^−1^s^−1^, correspondingly), have previously been identified as type II JM displacers, further supporting this binding mode.^73^ Additionally, in the active non-autoinhibited conformation of KIT, type II inhibitor ripretinib mimics the autoinhibitory lock by positioning its phenyl ring where W557 from the JM domain typically resides, stabilizing the DFG-out conformation (**Figure 9C**).^32^ In line with these structural insights, our results showed that ripretinib exhibited the slowest association (6.2 M^−1^s^−1^) and dissociation rates (1.4-day residence time) in the autoinhibited conformation, reflecting the substantial conformational adaptation required for binding.

In this context, various studies have demonstrated that kinase inhibitors, by interacting with specific kinase conformations, can suppress kinase activity while increasing protein expression, promoting ubiquitination and degradation, or facilitating protein-protein interactions that amplify non-canonical signaling pathways. For instance, the inhibitor BVD523 enhances the ubiquitination and degradation of active ERK2, amplifying its therapeutic potential, whereas some B-RAF inhibitors paradoxically activate signaling by promoting RAF dimerization.^74^ In the case of KIT, different studies have identified that certain inhibitors can induce KIT internalization and degradation. Under physiological conditions, KIT undergoes rapid internalization upon ligand engagement. Unexpectedly, imatinib and masitinib promote substantial internalization of the autoinhibited receptor in the absence of SCF (~50% at 2 μM). Likewise, GSK2606414 induces 50% downregulation of KIT at 250 nM, whereas dasatinib produces no detectable effect. ^74,75^

Strikingly, the kinetic profiles of these inhibitors correlate with their ability to induce KIT degradation. Back-cleft binding type II inhibitors that promote degradation—imatinib (*k*_*on*_ = 24 M^−1^s^−1^), masitinib (*k*_*on*_ = 38 M^−1^s^−1^), and GSK2606414 (*k*_*on*_ = 800 M^−1^s^−1^)—exhibit markedly slow association rates with autoinhibited KIT compared with the substantially faster binding observed for the non-autoinhibited form (**Table 1**). These slow *k*_*on*_ values reflect a significant kinetic barrier to binding, consistent with the need to displace the JM domain from its autoinhibitory position to gain access to the back cleft. In contrast, dasatinib, which binds the gate area shows rapid association with both autoinhibited (*k*_*on*_ = 8.7×10^4^ M^−1^s^−1^) and non-autoinhibited KIT (*k*_*on*_ = 1.2×10^6^ M^−1^s^−1^) and does not induce significant KIT degradation. Together, the kinetic signatures observed for these “kinase degraders” suggest that the displacement of the JM domain in autoinhibited KIT could potentially lead to a conformation that cells might recognize as similar to the active, non-autoinhibited form of KIT, thereby triggering internalization and degradation pathways.

### Avapritinib e ploits inetic selectivity to achieve durable KIT D816V inhibition with transient off-target activity

Although selectivity is often assessed through equilibrium-based parameters such as IC_50_ or *K*_*d*_, the role of kinetic selectivity remains comparatively underexplored. Unlike affinity metrics, which offer a static snapshot of drug-target binding, kinetic parameters capture the dynamic nature of molecular interactions over time. Drug selectivity is not static, it evolves over the dosing interval depending on how long a compound engages both its intended and unintended targets. Importantly, kinetic and affinity selectivity do not always correlate. A drug may exhibit little affinity discrimination between targets yet display substantial kinetic selectivity by maintaining long residence time on its main target while dissociating rapidly from collateral targets. This kinetic selectivity may maximize the therapeutic window, whereas sustained binding to toxicity-associated targets may compromise safety and narrow the therapeutic window.^10–12^

To study the effect of residence time on avapritinib and sunitinib’s selectivity profile, we characterized these compounds in parallel with 141 different kinases (**Figure 11, Supplementary spreadsheet-Kinetic Selectivity**). Based on affinity alone, both compounds displayed selectivity patterns consistent with prior published data. ^64^ Avapritinib bound with high affinity (*K*_*d*_ ≤ 100 nM) to 8% of the kinases tested, while sunitinib showed broad activity, interacting with 45% of them. In agreement with prior cellular studies, avapritinib also demonstrated reduced activity against the structurally related FLT3 D835Y mutant and FLT3 ITD.^76^ However, when residence time was taken into account, avapritinib exhibited pronounced kinetic selectivity: it dissociated rapidly (within seconds to 1.8 min) from off-targets but remained bound to its primary target, KIT D816V, for approximately one hour. This kinetic selectivity likely contributes to its manageable toxicity in clinical trials and high response rates among patients with KIT D816V-driven disease.^4^

**Figure 11:**
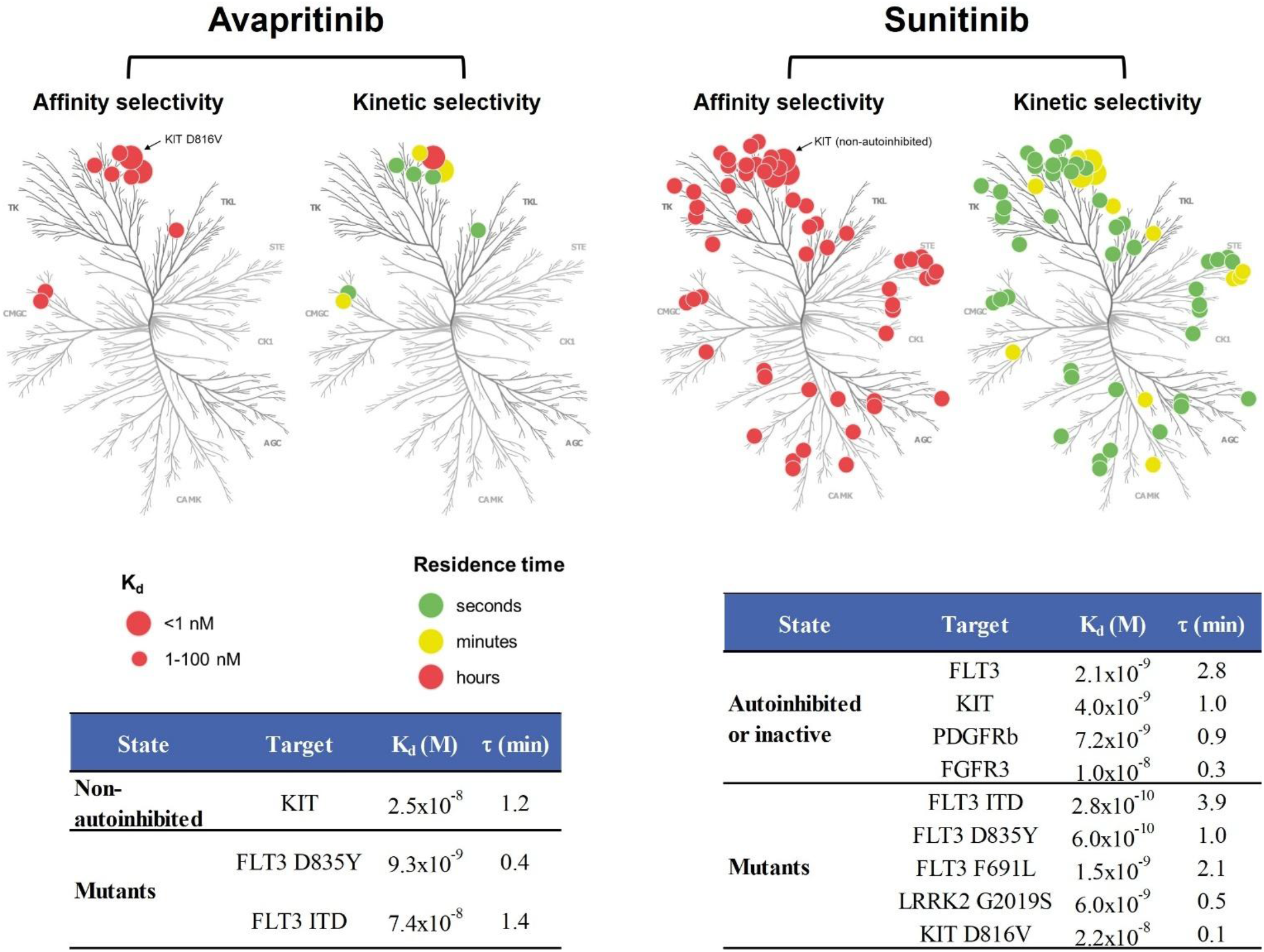
Kinome response to avapritinib and sunitinib using KINETICfinder. Left panels illustrate the kinome drug sensitivity against 141 kinases based on *K*_*d*_ values represented by red circles, where size indicates affinity magnitude. Right panels depict kinetic selectivity based on residence time, with circle colors reflecting the magnitude of this parameter. Tables list affinity and residence time (τ) values for those targets not shown in the kinome tree.

The distinct kinome selectivity profiles of avapritinib and sunitinib are reflected in their divergent adverse event (AEs) patterns. While the most reported AEs with avapritinib (nausea, diarrhea, anemia) are shared with other KIT inhibitors such as imatinib and sunitinib, toxicities frequently associated with broader-spectrum inhibitors, such as sunitinib (skin discoloration, severe myelosuppression, hypertension, and hypothyroidism), were uncommon or not observed with avapritinib.^77,78^ However, avapritinib exhibits a higher incidence of neuropsychiatric AEs compared to other KIT inhibitors. These effects likely arise from its ability to penetrate the blood–brain barrier (BBB) and subsequent interaction with unidentified CNS targets. Similar cognitive AEs have been reported with other BBB-penetrant kinase inhibitors (entrectinib and lorlatinib) that do not inhibit KIT or PDGFRα.^64,77^

### FF-1 1 1 covalently targets the non-autoinhibited conformation of KIT

Covalent inhibitors have proven highly effective in clinical cancer therapy, offering distinct pharmacological advantages over reversible compounds. Despite recent successes in the rational design of covalent KIT inhibitors, many rational strategies have failed, and to date, only a single covalent KIT inhibitor has been structurally confirmed by X-ray crystallography (PDB 6xVB). Moreover, just 22% of FDA-approved covalent drugs were discovered through rational approaches, underscoring the need for innovative strategies to identify covalent molecules—a challenge particularly relevant for overcoming the limitations of reversible inhibitors in patients with advanced GIST.^12,13^

Irreversible inhibition generally proceeds through a two-step mechanism, beginning with an initial reversible binding event, followed by a time-dependent inactivation driven by covalent bond formation. We therefore hypothesized that this kinetic signature, manifested as time-dependent inhibition, could serve as a marker for covalent engagement of the target. To explore this, we screened 23 irreversible inhibitors against KIT and identified three compounds—FF-10101, 5-7-Oxozeaenol, and FIIN-2—that exhibited both strong affinity and pronounced time-dependent inhibition. Interestingly, these inhibitors were originally designed to covalently target FLT3, MKK7, and FGFR1-4, respectively. The observation that they also inhibit KIT in a time-dependent manner suggests that they may engage the receptor through covalent modification.

To validate this hypothesis and gain insight into the binding mode, we selected FF-10101 for detailed analysis (**Figure 12**). Non-autoinhibited KIT was preincubated with either DMSO (left) or FF-10101 (right), followed by tryptic digestion and LC-MS/MS peptide mapping to identify the site of modification. This approach revealed the peptide TC_FF-10101_WDADPLK exclusively in the FF-10101-treated sample. We further confirmed this modification using parallel reaction monitoring (PRM), a high-resolution, high-accuracy technique that enabled unambiguous tracking of both the unmodified peptide (TCWDADPLKK, left) and its Cys-modified counterpart (TC_FF-10101_WDADPLKK, right) across the elution profile. As shown in **Figure 12A-B**, while the unmodified peptide is present under both conditions, the covalently modified peptide appears only following FF-10101 treatment (bottom), with no detectable signal in the DMSO control (top).

**Figure 12.**
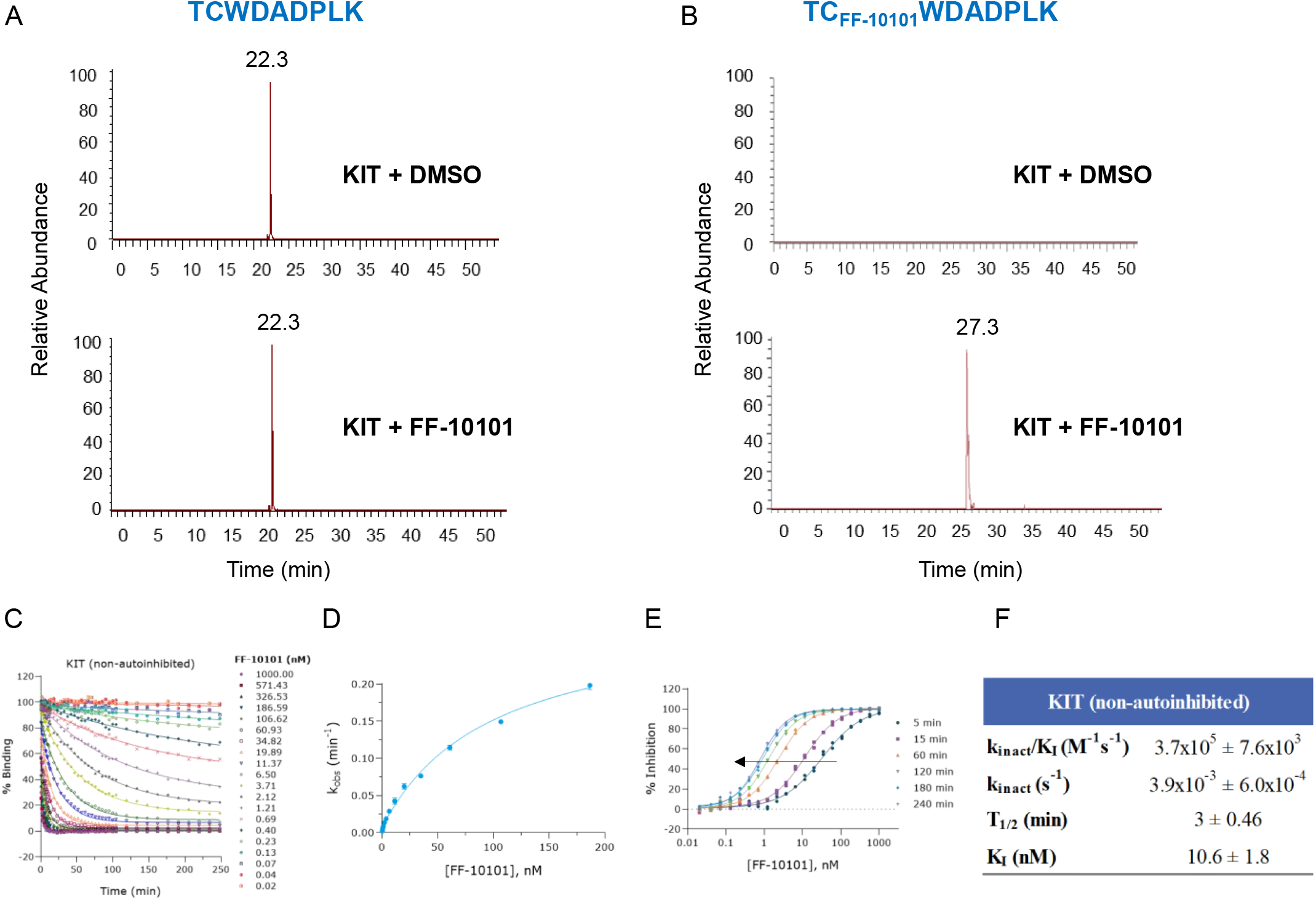
Biochemical evaluation of FF-10101. MS^2^ extracted ion chromatogram of the non-modified (A) and FF-10101-modified (B) peptide obtained in the parallel reaction monitoring analysis of the non-autoinhibited KIT kinase domain treated with DMSO (Top) or FF-10101 (Bottom). COVALfinder kinetic assays for (C-F) the non-autoinhibited KIT. Plots show progress curves, dependence of *k*_*obs*_ on compound concentration, dose-response curves over time and inactivation constants. Values represent mean ± SD from two independent experiments.

Complementary kinetic analysis using the COVALfinder^®^ TR-FRET platform corroborated these findings (**Figure 12 C-F**), demonstrating that FF-10101 irreversibly engages the non-autoinhibited form of KIT. The interaction is characterized by a rapid inactivation rate (3.9×10^−3^ s^−1^), low *K*_*I*_ (10.6 nM) and relatively high inactivation efficiency (3.7×10^5^ M^−1^s^−1^).

## Conclusions

Through comprehensive kinetic profiling of 172 kinase inhibitors across the autoinhibited, non-autoinhibited and D816V-mutated states of KIT, our study revealed that distinct conformations demand different requirements for kinetic optimization. Inhibitors targeting KIT D816V, characterized by a destabilized A-loop and increased structural plasticity, achieved high efficacy primarily through prolonged residence time. In contrast, the non-autoinhibited state presented a more complex optimization landscape: inhibitors stabilizing the DFG-in conformation required extended residence times similar to D816V inhibitors, whereas those targeting DFG-out conformations benefited most from accelerated association rates, with residence time playing a secondary role.

Systematic kinetic profiling also allowed us to distinguish kinetic fingerprints with direct therapeutic relevance. Prolonged residence time emerged as a hallmark of therapeutic success for both wild-type KIT and the D816V mutation, particularly in FDA-approved drugs such as imatinib, ripretinib, or ponatinib. Rather than relying solely on affinity, these compounds appear to achieve efficacy by sustaining target engagement long enough to outlast receptor turnover and prevent reactivation of downstream signaling. Avapritinib and midostaurin, both clinically effective against KIT D816V, displayed a similar kinetic signature characterized by markedly prolonged residence time. Given the D816V-associated rearrangement of the G-loop and αC-helix, it is plausible that inhibitors capable of engaging the FP-II region or extending into the adjacent Gα-pocket gain access to an expanded or more complementary binding interface. Consistent with this interpretation, other FP-II binders share this characteristic, pointing to a shared mechanism of kinetic stabilization via G-loop engagement.

These analyses further demonstrate that KIT binding kinetics are shaped by activation-linked structural features. Inhibitors stabilizing DFG-out conformations or accessing the back cleft consistently showed prolonged residence times, whereas αC-helix-in and front-cleft binders remained short-lived. R-spine engagement also influenced kinetic behavior, with inhibitors contacting two or more R-spine residues showing markedly extended residence times specifically against autoinhibited and non-autoinhibited KIT. Collectively, these findings emphasize that stabilizing the DFG-out or αC-helix out conformation, engaging the back pocket, and interacting with the R-spine are key strategies for achieving sustained KIT inhibition.

In the context of KIT D816V, our study points to faster dissociation as a mechanism of therapy resistance, underscoring the value of *k*_*off*_ in guiding inhibitor design. Beyond efficacy and resistance, our findings highlight the biological relevance of kinetic signatures in KIT regulation. We speculate that inhibitors that displace the JM domain from its autoinhibitory position may induce conformational states resembling receptor activation, thereby promoting KIT internalization and degradation. This raises the intriguing possibility that certain kinetic fingerprints could serve as indicators of “kinase degraders”, offering a novel strategy for selective protein downregulation. Additionally, kinetic profiling demonstrated its capability to uncover unexpected covalent interactions, as illustrated by FF-10101. Despite being developed for FLT3, FF-10101 formed irreversible adducts with KIT, an observation made possible through detection of time-dependent inhibition kinetics and mass spectrometry validation.

Finally, kinetic profiling provides an additional tool to evaluate and optimize kinase inhibitors for more selective engagement and potentially safer treatment options. Unlike static affinity measurements, kinetic selectivity captures the temporal dynamics of drug-target interactions, offering insights into differential engagement across kinases. Avapritinib exemplifies the potential and boundaries of kinetic selectivity as a tool to guide drug optimization. It displays remarkable kinetic discrimination, with prolonged residence on KIT D816V and rapid dissociation from collateral kinases, which likely contributes to its clinical efficacy and favorable safety profile. Nonetheless, avapritinib shows a higher incidence of neuropsychiatric AEs relative to other KIT inhibitors, reflecting its ability to cross the blood–brain barrier and interact with CNS targets not captured within our kinase panel. This underscores that, while kinetic profiling is a powerful approach for evaluating and optimizing target selectivity, complementary model systems are essential to detect off-target liabilities that extend beyond the profiled kinome.

While centered on KIT, this study offers a compelling argument for rethinking how drug performance is assessed— moving beyond static affinity metrics toward the dynamic framework of binding kinetics. Kinetic fingerprints uncover mechanistic and clinically meaningful features invisible at equilibrium, offering a richer understanding of efficacy, resistance, selectivity and mechanisms of drug action. Importantly, this conceptual and experimental framework is not limited to receptor tyrosine kinases. The intrinsic conformational plasticity of many proteins, coupled with structural perturbations induced by disease-associated mutations, makes this approach broadly applicable to diverse therapeutic targets, including non-RTKs, GPCRs, ion channels, epigenetic regulators or proteases. By capturing the temporal dimension of drug-target interactions, this methodology holds the potential to redefine how compounds are evaluated and prioritized, ultimately improving clinical success and guiding the rational design of next-generation therapeutics.

## Experimental Section

All reagents and experimental protocols used in this study are described in detail in the Supporting Information.

## Supporting Information

Experimental procedures, supplementary figures with phosphopeptide MS^2^ spectra and kinetic data as well as supplementary tables of FDA-approved kinase inhibitors and binding modes of kinase inhibitors have been included in the Supporting Information Word document. Results from kinetic profiling of compounds and kinetic selectivity of avapritinib and sunitinib have been included in separate Excel files.

## Acknowledgements

This study was supported by the Centre for the Development of Industrial Technology (CDTI) through the Horizon Europe EIC Accelerator Seal of Excellence SME grant (SoE-20211014). SK is grateful for support from the Structural Genomics Consortium (SGC), a registered charity (no: 1097737) that receives funds from Bristol Myers Squibb, Genome Canada through Ontario Genomics Institute, EU/EFPIA/OICR/McGill/KTH/Diamond Innovative Medicines Initiative 2 Joint Undertaking [EUbOPEN grant 875510], Janssen, Pfizer, and Takeda. SK is also funded by the German Cancer Research Center (DKTK), the Frankfurt Cancer Institute (FCI) and by the German Cancer Aid (Krebshilfe) pre-clinical drug development program TACTIC.

## Notes

### Competing Interest Statement

The authors have declared no competing interest.

## References

[1] Evans EK, Gardino AK, Kim JL, Hodous BL, Shutes A, Davis A, hu XJ, Schmidt-Kittler O, Wilson D, Wilson K, DiPietro L, hang Y, Brooijmans N, LaBranche TP, Wozniak A, Gebreyohannes YK, Schöffski P, Heinrich MC, DeAngelo DJ, Miller S, Wolf B, Kohl N, Guzi T, Lydon N, Boral A, Lengauer C. A precision therapy against cancers driven by KIT/PDGFRA mutations. Sci Transl Med. 2017 Nov 1;9(414):eaao1690.

[2] Laine E, Auclair C, Tchertanov L. Allosteric communication across the native and mutated KIT receptor tyrosine kinase. PLoS Comput Biol. 2012;8(8):e1002661.

[3] Klug LR, Kent JD, Heinrich MC. Structural and clinical consequences of activation loop mutations in class III receptor tyrosine kinases. Pharmacol Ther. 2018 Nov;191:123–134.

[4] Gilreath JA, Tchertanov L, Deininger MW. Novel approaches to treating advanced systemic mastocytosis. Clin Pharmacol. 2019 Jul 10;11:77–92.

[5] Agarwal S, Kazi JU, Rönnstrand L. Phosphorylation of the activation loop tyrosine 823 in c-Kit is crucial for cell survival and proliferation. J Biol Chem. 2013 Aug 2;288(31):22460–8.

[6] Sheikh E, Tran T, Vranic S, Levy A, Bonfil RD. Role and significance of c-KIT receptor tyrosine kinase in cancer: A review. Bosn J Basic Med Sci. 2022 Sep 16;22(5):683–698.

[7] Abdellateif MS, Bayoumi AK, Mohammed MA. c-Kit Receptors as a Therapeutic Target in Cancer: Current Insights. Onco Targets Ther. 2023 Sep 27;16:785–799.

[8] Mühlenberg T, Falkenhorst J, Schulz T, Fletcher BS, Teuber A, Krzeciesa D, Klooster I, Lundberg M, Wilson L, Lategahn J, von Mehren M, Grunewald S, Tüns AI, Wardelmann E, Sicklick JK, Brahmi M, Serrano C, Schildhaus HU, Sievers S, Treckmann J, Heinrich MC, Raut CP, Ou WB, Marino-Enriquez A, George S, Rauh D, Fletcher JA, Bauer S. KIT ATP-Binding Pocket/Activation Loop Mutations in GI Stromal Tumor: Emerging Mechanisms of Kinase Inhibitor Escape. J Clin Oncol. 2024 Apr 20;42(12):1439–1449.

[9] Gajiwala KS, Wu JC, Christensen J, Deshmukh GD, Diehl W, DiNitto JP, English JM, Greig MJ, He YA, Jacques SL, Lunney EA, McTigue M, Molina D, Quenzer T, Wells PA, Yu X, hang Y, ou A, Emmett MR, Marshall AG, hang HM, Demetri GD. KIT kinase mutants show unique mechanisms of drug resistance to imatinib and sunitinib in gastrointestinal stromal tumor patients. Proc Natl Acad Sci U S A. 2009 Feb 3;106(5):1542–7.

[10] Tavares FM, Gomes AC, Assunção EM, de Medeiros JLS, Scotti MT, Scotti L, Ishiki HM. Virtual Screening and Molecular Docking: Discovering Novel c-KIT Inhibitors. Curr Med Chem. 2022;29(2):166–188.

[11] Godesi S, Lee J, Nada H, Quan G, Elkamhawy A, Choi Y, Lee K. Small Molecule c-KIT Inhibitors for the Treatment of Gastrointestinal Stromal Tumors: A Review on Synthesis, Design Strategies, and Structure-Activity Relationship (SAR). Int J Mol Sci. 2023 May 29;24(11):9450.

[12] McAulay K, Hoyt EA, Thomas M, Schimpl M, Bodnarchuk MS, Lewis HJ, Barratt D, Bhavsar D, Robinson DM, Deery MJ, Ogg DJ, Bernardes GJL, Ward RA, Waring MJ, Kettle JG. Alkynyl Benzoxazines and Dihydroquinazolines as Cysteine Targeting Covalent Warheads and Their Application in Identification of Selective Irreversible Kinase Inhibitors. J Am Chem Soc. 2020 Jun 10;142(23):10358–10372.

[13] Schulz T, Gontla R, Teuber A, Beerbaum M, Fletcher BS, Mühlenberg T, Kaitsiotou H, Hardick J, Jeyakumar K, Keul M, Müller MP, Sievers S, Bauer S, Rauh D. Design, Synthesis, and SAR of Covalent KIT and PDGFRA Inhibitors─Exploring Their Potential in Targeting GIST. J Med Chem. 2025 Feb 13;68(3):3238–3259.

[14] Copeland RA, Pompliano DL, Meek TD. Drug-target residence time and its implications for lead optimization. Nat Rev Drug Discov. 2006 Sep;5(9):730–9.

[15] Copeland RA. Evolution of the drug-target residence time model. Expert Opin Drug Discov. 2021 Dec;16(12):1441–1451.

[16] Liu H, hang H, IJzerman AP, Guo D. The translational value of ligand-receptor binding kinetics in drug discovery. Br J Pharmacol. 2024 Nov;181(21):4117–4129.

[17] Georgi V, Andres D, Fernandez-Montalvan AE, Stegmann CM, Becker A, Mueller-Fahrnow A. Binding kinetics in drug discovery - A current perspective. Front Biosci (Landmark Ed). 2017 Jan 1;22(1):21–47.

[18] Lindberg MF, Deau E, Arfwedson J, George N, George P, Alfonso P, Corrionero A, Meijer L. Comparative Efficacy and Selectivity of Pharmacological Inhibitors of DYRK and CLK Protein Kinases. J Med Chem. 2023 Mar 23;66(6):4106–4130.

[19] Roy MJ, Winkler S, Hughes SJ, Whitworth C, Galant M, Farnaby W, Rumpel K, Ciulli A. SPR-Measured Dissociation Kinetics of PROTAC Ternary Complexes Influence Target Degradation Rate. ACS Chem Biol. 2019 Mar 15;14(3):361–368.

[20] Riching KM, Caine EA, Urh M, Daniels DL. The importance of cellular degradation kinetics for understanding mechanisms in targeted protein degradation. Chem Soc Rev. 2022 Jul 18;51(14):6210–6221.

[21] Mabanglo MF, Wilson B, Noureldin M, Kimani SW, Mamai A, Krausser C, González-Álvarez H, Srivastava S, Mohammed M, Hoffer L, Chan M, Avrumutsoae J, Li ASM, Hajian T, Tucker S, Green S, Szewczyk M, Barsyte-Lovejoy D, Santhakumar V, Ackloo S, Loppnau P, Li Y, Seitova A, Kiyota T, Wang JG, Privé GG, Kuntz DA, Patel B, Rathod V, Vala A, Rout B, Aman A, Poda G, Uehling D, Ramnauth J, Halabelian L, Marcellus R, Al-Awar R, Vedadi M. Crystal structures of DCAF1-PROTAC-WDR5 ternary complexes provide insight into DCAF1 substrate specificity. Nat Commun. 2024 Nov 23;15(1):10165.

[22] Imaide S, Riching KM, Makukhin N, Vetma V, Whitworth C, Hughes SJ, Trainor N, Mahan SD, Murphy N, Cowan AD, Chan KH, Craigon C, Testa A, Maniaci C, Urh M, Daniels DL, Ciulli A. Trivalent PROTACs enhance protein degradation via combined avidity and cooperativity. Nat Chem Biol. 2021 Nov;17(11):1157–1167.

[23] Wilms G, Schofield K, Maddern S, Foley C, Shaw Y, Smith B, Basantes LE, Schwandt K, Babendreyer A, Chavez T, McKee N, Gokhale V, Kallabis S, Meissner F, Rokey SN, Dunckley T, Montfort WR, Becker W, Hulme C. Discovery and Functional Characterization of a Potent, Selective, and Metabolically Stable PROTAC of the Protein Kinases DYRK1A and DYRK1B. J Med Chem. 2024 Oct 10;67(19):17259–17289.

[24] Daryaee F, hang Gogarty KR, Li Y, Merino J, Fisher SL, Tonge PJ. A quantitative mechanistic PK/PD model directly connects Btk target engagement and in vivo efficacy. Chem Sci. 2017 May 1;8(5):3434–3443.

[25] Walkup GK, You Z, Ross PL, Allen EK, Daryaee F, Hale MR et al. Translating slow-binding inhibition kinetics into cellular and in vivo effects. Nat Chem Biol. 2015 Jun;11(6):416–23.

[26] de Witte WEA, Rottschäfer V, Danhof M, van der Graaf PH, Peletier LA, de Lange ECM. Modelling the delay between pharmacokinetics and EEG effects of morphine in rats: binding kinetic versus effect compartment models. J Pharmacokinet Pharmacodyn. 2018 Aug;45(4):621–635.

[27] Ramsey SJ, Attkins NJ, Fish R, van der Graaf PH. Quantitative pharmacological analysis of antagonist binding kinetics at CRF1 receptors in vitro and in vivo. Br J Pharmacol. 2011 Oct;164(3):992–1007.

[28] Lee KSS, Yang J, Niu J, Ng CJ, Wagner KM, Dong H, Kodani SD, Wan D, Morisseau C, Hammock BD. Drug-Target Residence Time Affects in Vivo Target Occupancy through Multiple Pathways. ACS Cent Sci. 2019 Sep 25;5(9):1614–1624.

[29] Corrionero A., Alfonso AP. Method for determining kinetic profiles in drug discovery. US 10, 261, 079 B2. 2019.

[30] Georgi V, Schiele F, Berger BT, Steffen A, Marin apata PA, Briem H, Menz S, Preusse C, Vasta JD, Robers MB, Brands M, Knapp S, Fernández-Montalván A. Binding Kinetics Survey of the Drugged Kinome. J Am Chem Soc. 2018 Nov 21;140(46):15774–15782.

[31] Davis MI, Hunt JP, Herrgard S, Ciceri P, Wodicka LM, Pallares G, Hocker M, Treiber DK, arrinkar PP. Comprehensive analysis of kinase inhibitor selectivity. Nat Biotechnol. 2011 Oct 30;29(11):1046–51.

[32] Smith BD, Kaufman MD, Lu WP, Gupta A, Leary CB, Wise SC, Rutkoski TJ, Ahn YM, Al-Ani G, Bulfer SL, Caldwell TM, Chun L, Ensinger CL, Hood MM, McKinley A, Patt WC, Ruiz-Soto R, Su Y, Telikepalli H, Town A, Turner BA, Vogeti L, Vogeti S, Yates K, Janku F, Abdul Razak AR, Rosen O, Heinrich MC, Flynn DL. Ripretinib (DCC-2618) Is a Switch Control Kinase Inhibitor of a Broad Spectrum of Oncogenic and Drug-Resistant KIT and PDGFRA Variants. Cancer Cell. 2019 May 13;35(5):738–751.e9.

[33] Roskoski R Jr. Properties of FDA-approved small molecule protein kinase inhibitors: A 2024 update. Pharmacol Res. 2024 Feb;200:107059.

[34] van Linden OP, Kooistra AJ, Leurs R, de Esch IJ, de Graaf C. KLIFS: a knowledge-based structural database to navigate kinase-ligand interaction space. J Med Chem. 2014 Jan 23;57(2):249–77.

[35] Li K, Wang M, Akoglu M, Pollard AC, Klecker JB, Alfonso P, Corrionero A, Prendiville N, Qu W, Parker MFL, Turkman N, Cohen JA, Tonge PJ. Synthesis and Preclinical Evaluation of a Novel Fluorine-18-Labeled Tracer for Positron Emission Tomography Imaging of Bruton’s Tyrosine Kinase. ACS Pharmacol Transl Sci. 2023 Feb 10;6(3):410–421.

[36] Shi X, Sousa LP, Mandel-Bausch EM, Tome F, Reshetnyak AV, Hadari Y, Schlessinger J, Lax I. Distinct cellular properties of oncogenic KIT receptor tyrosine kinase mutants enable alternative courses of cancer cell inhibition. Proc Natl Acad Sci U S A. 2016 Aug 16;113(33):E4784–93.

[37] Arter C, Trask L, Ward S, Yeoh S, Bayliss R. Structural features of the protein kinase domain and targeted binding by small-molecule inhibitors. J Biol Chem. 2022 Aug;298(8):102247.

[38] Roskoski R Jr. The role of small molecule Kit protein-tyrosine kinase inhibitors in the treatment of neoplastic disorders. Pharmacol Res. 2018 Jul;133:35–52.

[39] Lee PY, Yeoh Y, Low TY. A recent update on small-molecule kinase inhibitors for targeted cancer therapy and their therapeutic insights from mass spectrometry-based proteomic analysis. FEBS J. 2023 Jun;290(11):2845–2864.

[40] Modi V, Dunbrack RL Jr. Defining a new nomenclature for the structures of active and inactive kinases. Proc Natl Acad Sci U S A. 2019 Apr 2;116(14):6818–6827.

[41] Lake EW, Muretta JM, Thompson AR, Rasmussen DM, Majumdar A, Faber EB, Ruff EF, Thomas DD, Levinson NM. Quantitative conformational profiling of kinase inhibitors reveals origins of selectivity for Aurora kinase activation states. Proc Natl Acad Sci U S A. 2018 Dec 18;115(51):E11894–E11903.

[42] Singh AK, Sonawane P, Kumar A, Singh H, Naumovich V, Pathak P, Grishina M, Khalilullah H, Jaremko M, Emwas AH, Verma A, Kumar P. Challenges and Opportunities in the Crusade of BRAF Inhibitors: From 2002 to 2022. ACS Omega. 2023 Jul 26;8(31):27819–27844.

[43] Gehringer, M. (2020). Covalent Kinase Inhibitors: An Overview. In: Laufer, S. (eds) Protein kinase Inhibitors. Topics in Medicinal Chemistry, vol 36. Springer, Cham.

[44] Zhao Z, Wu H, Wang L, Liu Y, Knapp S, Liu Q, Gray NS. Exploration of type II binding mode: A privileged approach for kinase inhibitor focused drug discovery? ACS Chem Biol. 2014 Jun 20;9(6):1230–41.

[45] Zhang H, He F, Gao G, Lu S, Wei Q, Hu H, Wu Z, Fang M, Wang X. Approved Small-Molecule ATP-Competitive Kinases Drugs Containing Indole/Azaindole/Oxindole Scaffolds: R&D and Binding Patterns Profiling. Molecules. 2023 Jan 17;28(3):943.

[46] Wodicka LM, Ciceri P, Davis MI, Hunt JP, Floyd M, Salerno S, Hua XH, Ford JM, Armstrong RC, arrinkar PP, Treiber DK. Activation state-dependent binding of small molecule kinase inhibitors: structural insights from biochemistry. Chem Biol. 2010 Nov 24;17(11):1241–9.

[47] Vijayan RS, He P, Modi V, Duong-Ly KC, Ma H, Peterson JR, Dunbrack RL Jr, Levy RM. Conformational analysis of the DFG-out kinase motif and biochemical profiling of structurally validated type II inhibitors. J Med Chem. 2015 Jan 8;58(1):466–79.

[48] Sessa F, Villa F. Structure of Aurora B-INCENP in complex with barasertib reveals a potential transinhibitory mechanism. Acta Crystallogr F Struct Biol Commun. 2014 Mar;70(Pt 3):294–8.

[49] Martinez R III, Defnet A, Shapiro P. Avoiding or Co-Opting ATP Inhibition: Overview of Type III, IV, V, and VI Kinase Inhibitors. Next Generation Kinase Inhibitors. 2020 Jul 15:29–59.

[50] Li MJ, Wu G, Kaas Q, Jiang T, Yu RL. Development of efficient docking strategies and structure-activity relationship study of the c-Met type II inhibitors. J Mol Graph Model. 2017 Aug;75:241–249.

[51] Roskoski R Jr. The role of fibroblast growth factor receptor (FGFR) protein-tyrosine kinase inhibitors in the treatment of cancers including those of the urinary bladder. Pharmacol Res. 2020Jan;151:104567.

[52] Roskoski R Jr. Properties of FDA-approved small molecule phosphatidylinositol 3-kinase inhibitors prescribed for the treatment of malignancies. Pharmacol Res. 2021 Jun;168:105579.

[53] Qu L, Lin H, Dai S, Guo M, Chen X, Jiang L, hang H, Li M, Liang X Chen, Wei H, Chen Y. Structural insight into the macrocyclic inhibitor TPX-0022 of c-Met and c-Src. Comput Struct Biotechnol J. 2023 Nov 17;21:5712–5718.

[54] Roskoski R Jr. Janus kinase (JAK) inhibitors in the treatment of neoplastic and inflammatory disorders. Pharmacol Res. 2022 Sep;183:106362.

[55] Crawford JJ, Johnson AR, Misner DL, Belmont LD, Castanedo G, Choy R, Coraggio M, Dong L, Eigenbrot C, Erickson R, Ghilardi N, Hau J, Katewa A, Kohli PB, Lee W, Lubach JW, McKenzie BS, Ortwine DF, Schutt L, Tay S, Wei B, Reif K, Liu L, Wong H, Young WB. Discovery of GDC-0853: A Potent, Selective, and Noncovalent Bruton’s Tyrosine Kinase Inhibitor in Early Clinical Development. J Med Chem. 2018 Mar 22;61(6):2227–2245.

[56] Ayala-Aguilera CC, Valero T, Lorente-Macías Á, Baillache DJ, Croke S, Unciti-Broceta A. Small Molecule Kinase Inhibitor Drugs (1995-2021): Medical Indication, Pharmacology, and Synthesis. J Med Chem. 2022 Jan 27;65(2):1047–1131.

[57] Gao C, Grøtli M, Eriksson LA. Characterization of interactions and pharmacophore development for DFG-out inhibitors to RET tyrosine kinase. J Mol Model. 2015 Jul;21(7):167.

[58] Harada G, Drilon A. TRK inhibitor activity and resistance in TRK fusion-positive cancers in adults. Cancer Genet. 2022 Jun;264–265:33–39.

[59] Dai Y, Hartandi K, Ji Z, Ahmed AA, Albert DH, Bauch JL, Bouska JJ, Bousquet PF, Cunha GA, Glaser KB, Harris CM, Hickman D, Guo J, Li J, Marcotte PA, Marsh KC, Moskey MD, Martin RL, Olson AM, Osterling DJ, Pease LJ, Soni NB, Stewart KD, Stoll VS, Tapang P, Reuter DR, Davidsen SK, Michaelides MR. Discovery of N-(4-(3-amino-1H-indazol-4-yl)phenyl)-N’-(2-fluoro-5-methylphenyl)urea (ABT-869), a 3-aminoindazole-based orally active multitargeted receptor tyrosine kinase inhibitor. J Med Chem. 2007 Apr 5;50(7):1584–97.

[60] Hart S, Goh KC, Novotny-Diermayr V, Hu CY, Hentze H, Tan YC, Madan B, Amalini C, Loh YK, Ong LC, William AD, Lee A, Poulsen A, Jayaraman R, Ong KH, Ethirajulu K, Dymock BW, Wood JW. SB1518, a novel macrocyclic pyrimidine-based JAK2 inhibitor for the treatment of myeloid and lymphoid malignancies. Leukemia. 2011 Nov;25(11):1751–9.

[61] Shen Y, Chen X, He J, Liao D, zu X. Axl inhibitors as novel cancer therapeutic agents. Life Sci. 2018 Apr 1;198:99–111.

[62] Chiappa M, Petrella S, Damia G, Broggini M, Guffanti F, Ricci F. Present and Future Perspective on PLK1 Inhibition in Cancer Treatment. Front Oncol. 2022 Jun 2;12:903016.

[63] Mol CD, Dougan DR, Schneider TR, Skene RJ, Kraus ML, Scheibe DN, Snell GP, ou H, Sang BC, Wilson KP. Structural basis for the autoinhibition and STI-571 inhibition of c-Kit tyrosine kinase. J Biol Chem. 2004 Jul 23;279(30):31655–63.

[64] Teuber A, Schulz T, Fletcher BS, Gontla R, Mühlenberg T, ischinsky ML, Niggenaber J, Weisner J, Kleinbölting SB, Lategahn J, Sievers S, Müller MP, Bauer S, Rauh D. Avapritinib-based SAR studies unveil a binding pocket in KIT and PDGFRA. Nat Commun. 2024 Jan 2;15(1):63.

[65] Barlaam B, Anderton J, Ballard P, Bradbury RH, Hennequin LF, Hickinson DM, Kettle JG, Kirk G, Klinowska T, Lambert-van der Brempt C, Trigwell C, Vincent J, Ogilvie D. Discovery of A D8931, an Equipotent, Reversible Inhibitor of Signaling by EGFR, HER2, and HER3 Receptors. ACS Med Chem Lett. 2013 May 31;4(8):742–6.

[66] Mandel S, Hanke T, Mathea S, Chatterjee D, Saraswati H, Berger BT, Schwalm MP, Yamamoto S, Tawada M, Takagi T, Ahmed M, Röhm S, Corrionero A, Alfonso P, Baena M, Elson L, Menge A, Krämer A, Pereira R, Müller S, Krause DS, Knapp S. Repurposing of the RIPK1-Selective Benzo[1,4]oxazepin-4-one Scaffold for the Development of a Type III LIMK1/2 Inhibitor. ACS Chem Biol. 2025 May 16;20(5):1087–1098.

[67] Wentsch HK, Walter NM, Bührmann M, Mayer-Wrangowski S, Rauh D, aman GJR, Willemsen Seegers N, Buijsman RC, Henning M, Dauch D, ender L, Laufer S. Optimized Target Residence Time: Type I1/2 Inhibitors for p38α MAP Kinase with Improved Binding Kinetics through Direct Interaction with the R-Spine. Angew Chem Int Ed Engl. 2017 May 2;56(19):5363–5367.

[68] Bravo E Jr, Li Y, Lin DY, Srinivasan B, Barone M, Li SX, DelloRusso F, Rahiyanath AS, Corrionero A, Alfonso P, Prendiville N, Kozakov D, Andreotti AH, Tonge PJ. Modulating the Binding Kinetics of Bruton’s Tyrosine Kinase Inhibitors through Transition-State Effects. J Am Chem Soc. 2025 Aug 6;147(31):27876–27891.

[69] Lyczek A, Berger BT, Rangwala AM, Paung Y, Tom J, Philipose H, Guo J, Albanese SK, Robers MB, Knapp S, Chodera JD, Seeliger MA. Mutation in Abl kinase with altered drug-binding kinetics indicates a novel mechanism of imatinib resistance. Proc Natl Acad Sci U S A. 2021 Nov 16;118(46):e2111451118.

[70] Jin B, Ding K, Pan J. Ponatinib induces apoptosis in imatinib-resistant human mast cells by dephosphorylating mutant D816V KIT and silencing β-catenin signaling. Mol Cancer Ther. 2014 May;13(5):1217–30.

[71] hu X, Kim JL, Newcomb JR, Rose PE, Stover DR, Toledo LM, hao H, Morgenstern KA. Structural analysis of the lymphocyte-specific kinase Lck in complex with non-selective and Src family selective kinase inhibitors. Structure. 1999 Jun 15;7(6):651–61.

[72] Wagner AJ, Severson PL, Shields AF, Patnaik A, Chugh R, Tinoco G, Wu G, Nespi M, Lin J, hang Y, Ewing T, Habets G, Burton EA, Matusow B, Tsai J, Tsang G, Shellooe R, Carias H, Chan K, Rezaei H, Sanftner L, Marimuthu A, Spevak W, Ibrahim PN, Inokuchi K, Alcantar O, Michelson G, Tsiatis AC, hang C, Bollag G, Trent JC, Tap WD. Association of Combination of Conformation-Specific KIT Inhibitors With Clinical Benefit in Patients With Refractory Gastrointestinal Stromal Tumors: A Phase 1b/2a Nonrandomized Clinical Trial. JAMA Oncol. 2021 Sep 1;7(9):1343–1350.

[73] McTigue M, Murray BW, Chen JH, Deng YL, Solowiej J, Kania RS. Molecular conformations, interactions, and properties associated with drug efficiency and clinical performance among VEGFR TK inhibitors. Proc Natl Acad Sci U S A. 2012 Nov 6;109(45):18281–9.

[74] Jones LH. Small-Molecule Kinase Downregulators. Cell Chem Biol. 2018 Jan 18;25(1):30–35.

[75] Mahameed M, Wilhelm T, Darawshi O, Obiedat A, Tommy WS, Chintha C, Schubert T, Samali A, Chevet E, Eriksson LA, Huber M, Tirosh B. The unfolded protein response modulators GSK2606414 and KIRA6 are potent KIT inhibitors. Cell Death Dis. 2019 Apr 1;10(4):300.

[76] Weisberg E, Meng C, Case AE, Sattler M, Tiv HL, Gokhale PC, Buhrlage SJ, Liu X, Yang J, Wang J, Gray N, Stone RM, Adamia S, Dubreuil P, Letard S, Griffin JD. Comparison of effects of midostaurin, crenolanib, quizartinib, gilteritinib, sorafenib and BLU-285 on oncogenic mutants of KIT, CBL and FLT3 in haematological malignancies. Br J Haematol. 2019 Nov;187(4):488–501.

[77] Jones RL, Serrano C, von Mehren M, George S, Heinrich MC, Kang YK, Schöffski P, Cassier PA, Mir O, Chawla SP, Eskens FALM, Rutkowski P, Tap WD, hou T, Roche M, Bauer S. Avapritinib in unresectable or metastatic PDGFRA D842V-mutant gastrointestinal stromal tumours: Long-term efficacy and safety data from the NAVIGATOR phase I trial. Eur J Cancer. 2021 Mar;145:132–142.

[78] Aparicio-Gallego G, Blanco M, Figueroa A, García-Campelo R, Valladares-Ayerbes M, Grande-Pulido E, Antón-Aparicio L. New insights into molecular mechanisms of sunitinib-associated side effects. Mol Cancer Ther. 2011 Dec;10(12):2215–23.

[79] Bonzon-Kulichenko E, Pérez-Hernández D, Núñez E, Martínez-Acedo P, Navarro P, Trevisan-Herraz M, Ramos Mdel C, Sierra S, Martínez-Martínez S, Ruiz-Meana M, Miró-Casas E, García-Dorado D, Redondo JM, Burgos JS, Vázquez J. A robust method for quantitative high-throughput analysis of proteomes by 18O labeling. Mol Cell Proteomics. 2011 Jan;10(1):M110.003335.

[80] Lozano-Prieto M, Camafeita E, Jorge I, Laguillo-Gómez A, Barrero-Rodríguez R, Devesa CA, Pertusa C, Calvo E, Sánchez-Madrid F, Vázquez J, Martin-Cofreces NB. In-gel protein digestion using acidic methanol produces a highly selective methylation of glutamic acid residues. J Proteomics. 2024 Jul 30;304:105229.

[81] Martínez-Bartolomé S, Navarro P, Martín-Maroto F, López-Ferrer D, Ramos-Fernández A, Villar M, García-Ruiz JP, Vázquez J. Properties of average score distributions of SEQUEST: the probability ratio method. Mol Cell Proteomics. 2008 Jun;7(6):1135–45.

[82] Motulsky HJ, Mahan LC. The kinetics of competitive radioligand binding predicted by the law of mass action. Mol Pharmacol. 1984 Jan;25(1):1–9.

[83] Mandel S, Hanke T, Prendiville N, Baena-Nuevo M, Berger LM, Farges F, Schwalm MP, Berger BT, Kraemer A, Elson L, Saraswati H, Abdul Azeez KR, Dederer V, Mathea S, Corrionero A, Alfonso P, Keller S, Gstaiger M, Krause DS, Müller S, Röhm S, Knapp S. Covalent Targeting Leads to the Development of a LIMK1 Isoform-Selective Inhibitor. J Med Chem. 2025 Jul 2. doi: 10.1021/acs.jmedchem.5c01204. Epub ahead of print.

