## Supplementary information for "Kinetic Fingerprints as Mechanistic and Clinical Roadmaps Across KIT Activation States"

[b] Ana Corrionero.
Department of Biotechnology-Plant Biology, School of Agricultural, Food and Biosystems Engineering.
Universidad Politécnica de Madrid.
28040 Madrid, Spain.

[c] Sandra Röhm, Stefan Knapp.
Institute for Pharmaceutical Chemistry.
Johann Wolfgang Goethe-University.
Max-von-Laue-Str. 9, D-60438 Frankfurt am Main, Germany.

[d] Sandra Röhm, Stefan Knapp.
Structure Genomics Consortium Buchmann Institute for Molecular Life Sciences.
Johann Wolfgang Goethe-University.
Max-von-Laue-Str. 15, D-60438 Frankfurt am Main, Germany.

[e] Emilio Camafeita.
Cardiovascular Proteomics Laboratory.
Centro Nacional de Investigaciones Cardiovasculares (CNIC).
C/Melchor Fernández Almagro, 3, 28029 Madrid, Spain.

[f] Emilio Camafeita.
Centro de Investigación Biomédica en Red.
Enfermedades Cardiovasculares (CIBERCV).
Av. Monforte de Lemos, 3-5, 28029 Madrid, Spain.

### TABLE OF CONTENTS

**EXPERIMENTAL SECTION** 3

*Reagents* 3

*Sample Preparation for Proteomics Analysis* 3

*Shotgun Liquid Chromatography - Tandem Mass Spectrometry (LC-MS/MS) Analysis* 3

*High-Throughput Kinetic Screening Assay for Reversible Compounds* 3

*Kinetic Selectivity Assay* 3

*High-Throughput Kinetic Screening Assay for Irreversible Compounds* 4

*Confirmation of Covalent Binding of FF10101 to KIT by MS Analysis* 4

*Statistical Methods* 4

**SUPPLEMENTARY FIGURES** 5

*Figure S1. $MS^2$ fragmentation spectrum from peptides bearing phosphorylated tyrosine residues in the JM domain.* 5

*Figure S2. $MS^2$ fragmentation spectrum from precursor ions containing peptides bearing phosphorylated tyrosine residues in the kinase domain.* 6

*Figure S3. Comparison of $K_d$ (A), $k_{on}$ (B) and $k_{off}$ (C) values measured with KINETICfinder and those reported in the literature with KIT non autoinhibited and KIT D816V.* 7

*Figure S4. Comparison of association and dissociation rate constants with affinity values measured for non-autoinhibited KIT* 8

*Figure S5. Distribution of inhibitor binding kinetics and affinity across KIT activation states, stratified by R-spine engagement.* 8

*Figure S6: Superimposition of crystal structures highlighting the plasticity of the KIT kinase domain and diverse conformational states.* 9

**SUPPLEMENTARY TABLES** 10

*Table S1. Compounds screened in the HTS kinetic assays.* 10

*Table S2. FDA-approved kinase inhibitors, their main kinase targets and therapeutic indications.* 14

*Table S3. Binding modes of the kinase inhibitors analyzed in this study across relevant proteins.* 17

#### EXPERIMENTAL SECTION

##### Reagents

Human histidine-tagged KIT (aa 544-976) was purchased from Life Technologies. No special measures were taken to activate this kinase. Human GST-tagged KIT (aa 544-976) was purchased from ProQuinase. This kinase was activated in-vitro via autophosphorylation with ATP. Human GST-tagged KIT D816V (aa 544-976) was purchased from Life Technologies. This kinase was activated in-vitro via autophosphorylation with ATP. Kinase inhibitors (Table S1) were purchased from MedChemExpress, Sigma-Aldrich, SelleckChem, ApexBio and Santa Cruz Biotechnology.

##### Sample Preparation for Proteomics Analysis

The protein samples were digested using the one-step in-gel method.[70] Then the protein bands were visualized by Coomassie and digested overnight at 37ºC with trypsin (Promega). The resulting tryptic peptides were desalted in RP C-18 extraction cartridges (Oasis; Waters, Milford, MA USA), and vacuum-dried.

##### Shotgun Liquid Chromatography - Tandem Mass Spectrometry (LC-MS/MS) Analysis

For peptide mapping, the dried peptide samples were taken up in 0.1% (v/v) formic acid and analyzed by LC-MS/MS on an Ultimate 3000 nano-HPLC apparatus (Dionex, Sunnyvale, CA USA) coupled to a hybrid quadrupole-orbitrap mass spectrometer (Q Exactive HF, Thermo Scientific) The peptides were separated in a C-18 RP nano-column (Thermo Scientific) using a 200 nL/min flow. For shotgun analysis, a continuous gradient consisting of 8-28% B for 30 min (B: 90% acetonitrile, 0.1% formic acid) was used. The resulting $MS^2$ spectra were searched with Proteome Discoverer 2.5 (Thermo Scientific) against the UniProtKB/Swiss-Prot human database using the following parameters: trypsin digestion with 2 maximum missed cleavage sites; precursor and fragment mass tolerances of 800 ppm and 0.02 Da, respectively; variable modifications: Cys carbamidomethylation and FF10101 adduct formation; Met oxidation; Glu methylation[71]; and Ser, Thr, and Tyr phosphorylation. The false discovery rate (FDR) of peptide identification was calculated using the probability ratio method[72] after precursor post-filtering at 15 ppm. For targeted parallel reaction monitoring analysis, a continuous gradient consisting of 6-25% B for 15 min (B: 90% acetonitrile, 0.1% formic acid) was used. Each MS run consisted of enhanced FT-resolution spectra (120,000 resolution) in the 150–1,600 m/z range followed by data-independent acquisition of $MS^2$ spectra from precursor ions 789.8979 (+2) m/z and 553.2528 (+2) m/z, corresponding to the Cys-modified and non-modified TCFF-WDADPLK peptide, respectively. This data-independent MS and $MS^2$ scan cycle was repeated along the chromatographic run. XCalibur 2.2 (Thermo Scientific) was used to obtain the extracted ion chromatograms of selected ions from $MS^2$ scans.

##### High-Throughput Kinetic Screening Assay for Reversible Compounds

KINETICfinder assays were conducted in black 384-well microplates (Greiner) containing either 0.2 nM autoinhibited KIT, 0.1 nM non-autoinhibited KIT or 2 nM KIT D816V. The reaction mixture included 10 nM fluorescent probe, 2 nM terbium-labeled antibody (Life Technologies) and assay buffer (50 mM HEPES, pH 7.5, 10 mM $MgCl_2$, 0.01% Brij-35, 1 mM TCEP and 1% DMSO). Test compounds were prepared in DMSO as 100X concentrated solutions and diluted in a 4-point, 10-fold series. The kinetic assays were read continuously at RT in a PHERAstar FSX plate reader (BMG LABTECH) and the specific signals were fitted to the Motulsky-Mahan's equation.[73] The $K_d$, $k_{on}$, $k_{off}$ and residence time ($\tau$) of each test compound were calculated using KINPy® software (Enzymlogic). Throughout the assay, the Z-factor and S/B ratio remained consistently above 0.5 and 2, respectively, while the %CV of the controls did not exceed 10%, indicating high assay quality. Off-rates were set to > 0.5 $s^{-1}$ if values are above assay quantitation limit.

##### Kinetic Selectivity Assay

Avapritinib and sunitinib were screened against a panel of 141 kinase targets using the KINETICfinder assay platform as previously described. $K_d$, $k_{on}$, $k_{off}$ and residence time on primary and secondary kinase targets were determined in singlicate 4-point dilution series of compounds. Across all targets analyzed, the Z-factor and S/B ratio remained consistently above 0.4 and 2, respectively, while the %CV of controls remained below 10%

throughout the assay, confirming robust performance. Kinome trees were generated using Coral (http://phanstiel-lab.med.unc.edu/CORAL/).

**High-Throughput Kinetic Screening Assay for Irreversible Compounds**

COVALfinder assays (Enzymlogic) were conducted in black 384-well microplates containing 0.1 nM of non-autoinhibited KIT. The reaction mixture included 10 nM fluorescent probe, 2 nM labeled antibody and assay buffer. For all experiments, test compounds were prepared as 100X stock solutions and serially diluted in DMSO. Kinetic measurements were continuously recorded at RT and the specific signals were fitted to a single exponential equation to determine the kinetic constants and the mechanism of inactivation (one- or two-step).[74] The same data sets were used to produce concentration−response curves at each time point to facilitate the inspection of time dependency. Throughout the assay, the Z-factor and S/B ratio remained consistently above 0.5 and 2, respectively, while the %CV of the controls did not exceed 10%, indicating high assay quality.

**Confirmation of Covalent Binding of FF10101 to KIT by MS Analysis**

Non-autoinhibited KIT protein (0.55 µM) was incubated with 10 µM of FF-110101 in assay buffer (50 mM HEPES, pH 7.5, 10 mM $MgCl_2$, 1 mM TCEP, 0.01% Brij-35, and 1% DMSO) at RT for 2 h. Negative control samples were prepared by adding DMSO instead of FF-110101. The covalent reaction was stopped by adding an equal volume of sample buffer and boiling the samples for 5 min at 95ºC. The samples were then subjected to targeted parallel reaction monitoring LC-MS/MS analysis following the treatment described in the *Sample Preparation for Proteomics Analysis* subsection.

**Statistical Methods**

Statistical analysis was performed using GraphPad Prism. Spearman correlation calculations were utilized to compute correlation coefficients. Agreement between methods was evaluated by Bland-Altman analysis using % mean log differences. To test the differences among populations, Kruskal–Wallis one-way ANOVA on ranks for unpaired samples, with use of Dunn’s post hoc test. All statistical tests were two-sided and a P value of less than 0.05 was considered statistically significant.

#### SUPPLEMENTARY FIGURES

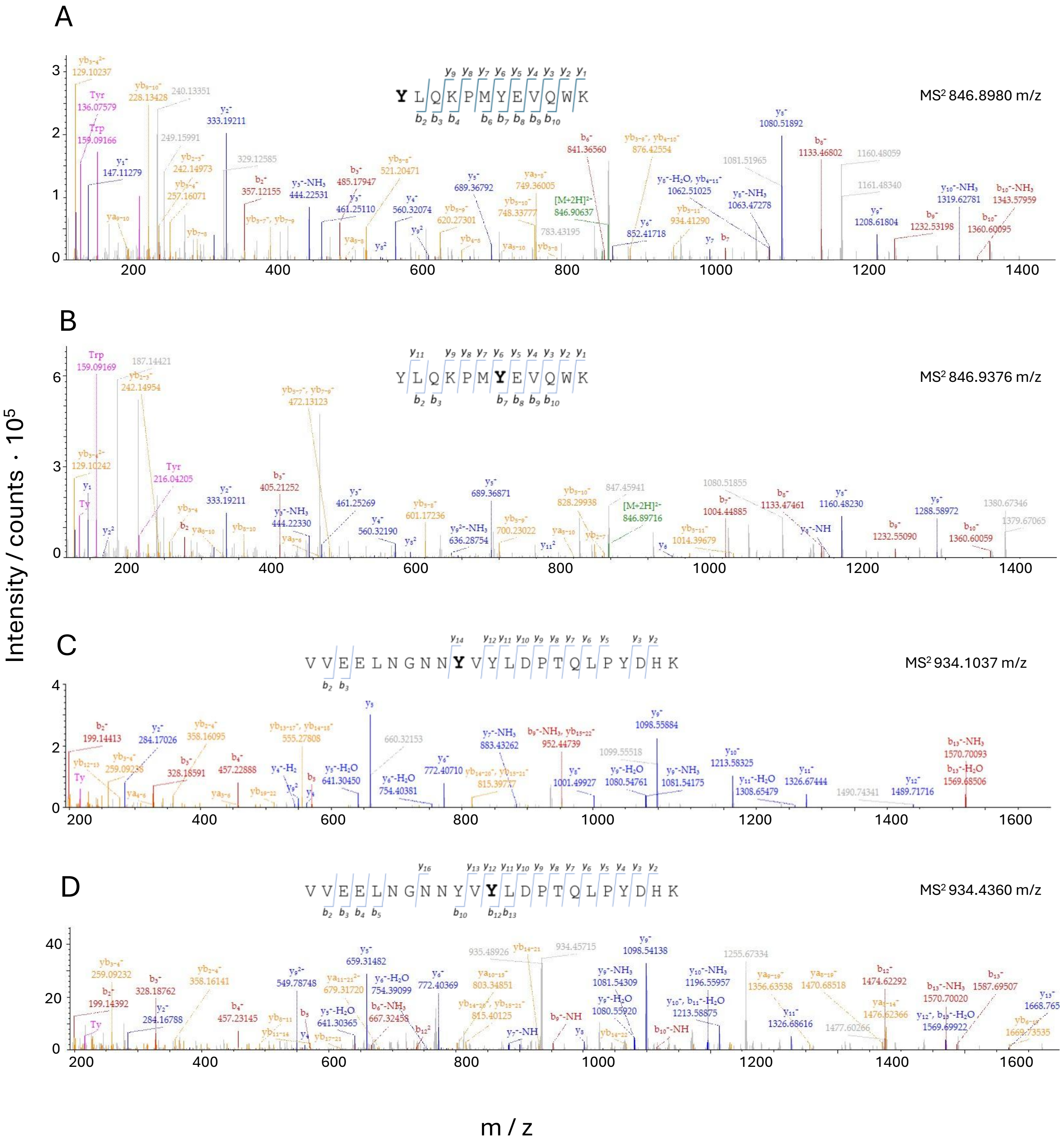


**Figure S1. MS$^2$ fragmentation spectrum from peptides bearing phosphorylated tyrosine residues in the JM domain.** Phosphorylation at Y547 (A), Y553 (B), Y568 (C) and Y570 (D) is observed exclusively in the non-autoinhibited form of the KIT receptor. Fragment ions are assigned to the main series: y-ions, containing the peptide C-terminus (blue), and b-ions, containing the N-terminus (red). Internal fragments, immonium ions and precursor ions are indicated in orange, pink, and green, respectively. Phosphorylated tyrosine residues are shown in bold within the corresponding peptide sequences.

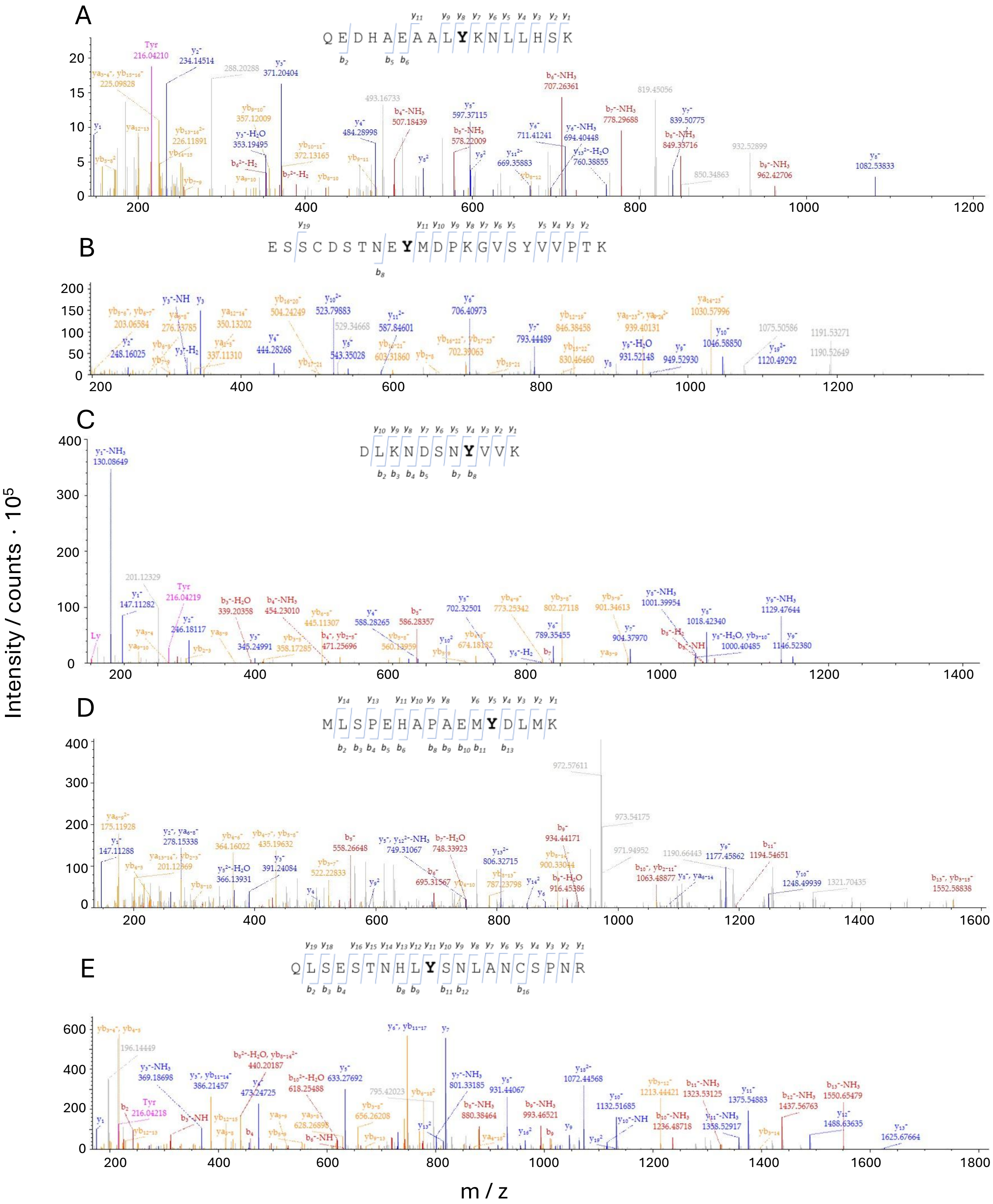


**Figure S2. MS² fragmentation spectrum from precursor ions containing peptides bearing phosphorylated tyrosine residues in the kinase domain.** Phosphorylation at (A) Y703, (B) Y721, (C) Y823, (D) Y900 and (E) Y936 is observed in the autoinhibited and non-autoinhibited forms of the KIT receptor. Fragment ions are assigned to the main series: y-ions, from containing the peptide C-terminus (blue), and b-ions, from containing the N-terminus (red). Internal fragments, immonium ions and precursor ions are indicated in orange, pink, and green, respectively. Phosphorylated tyrosine residues are shown in bold within the corresponding peptide sequences.

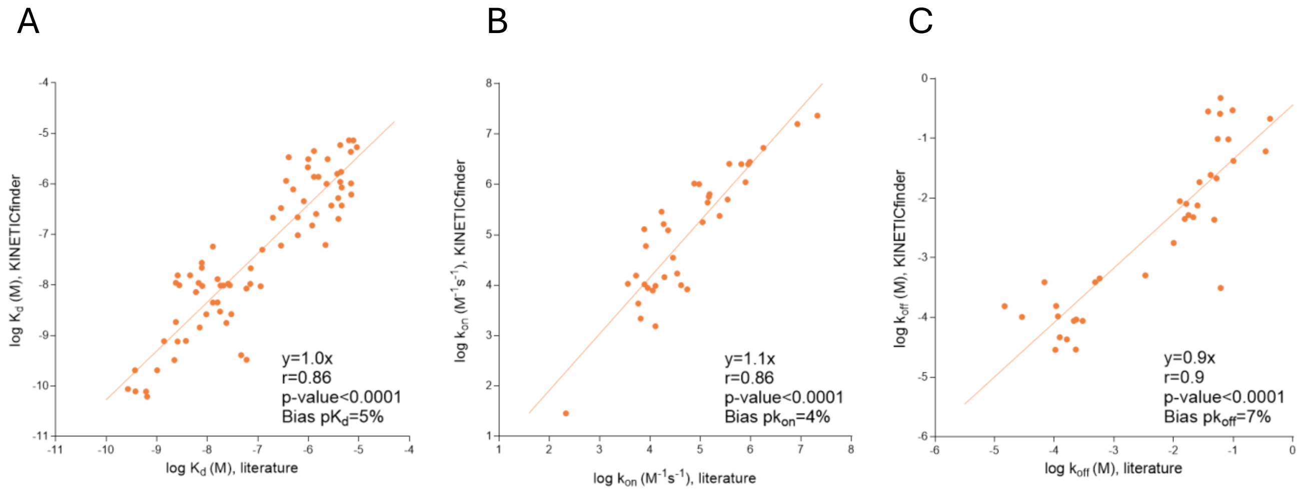


**Figure S3. Comparison of $K_d$ (A), $k_{on}$ (B) and $k_{off}$ (C) values measured with KINETICfinder and those reported in the literature with KIT non autoinhibited and KIT D816V.** Only parameters within the assay's detection limits are shown. Spearman correlation coefficients (r), p-values and and Bland–Altman statistics (bias) are provided in the graphs.

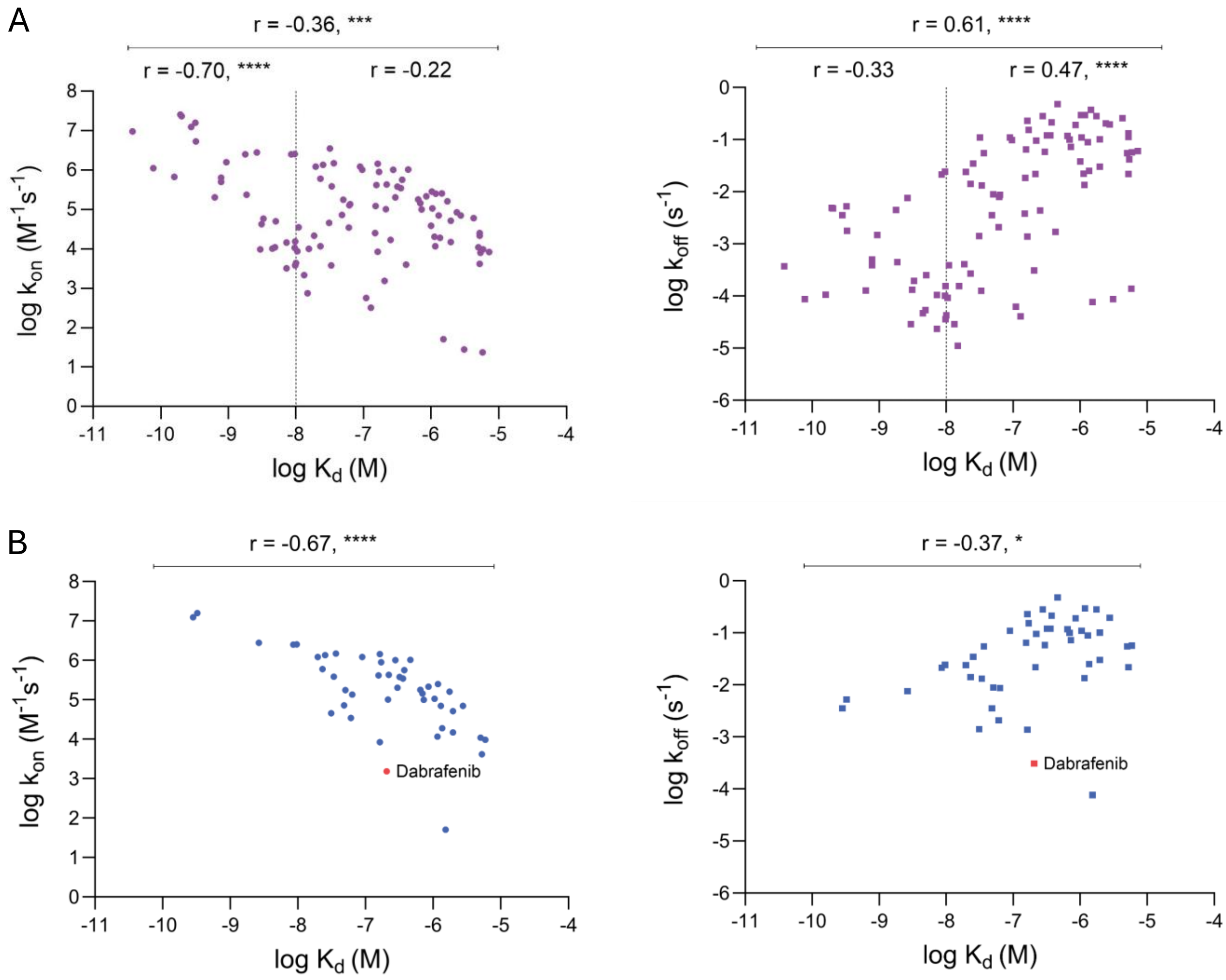

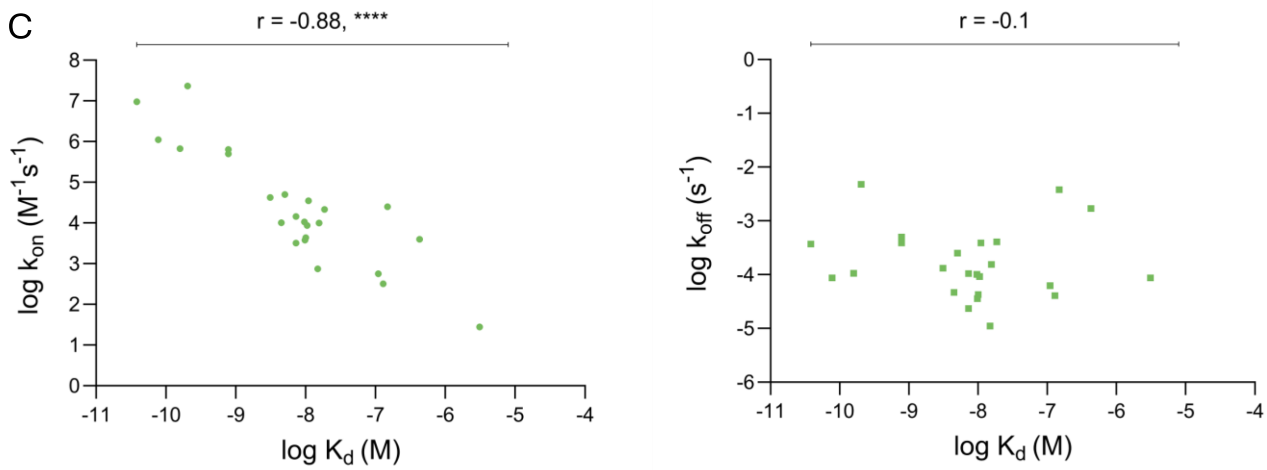


**Figure S4. Comparison of association and dissociation rate constants with affinity values measured for non-autoinhibited KIT.** (A) Data for all inhibitors tested with non-autoinhibited KIT. (B–C) Analysis of inhibitors annotated in the KLIFS database as targeting the (B) DFG-in or (C) DFG-out states of non-autoinhibited KIT, regardless of αC-helix orientation (in or out). Only parameters within the assay's detection limits are shown. Spearman correlation coefficients (r) and p-values for the entire dataset and two affinity groups (indicated by the lines) are provided above the graphs. *, $p < 0.05$; **, $p < 0.01$; ***, $p < 0.001$; ****, $p < 0.0001$.

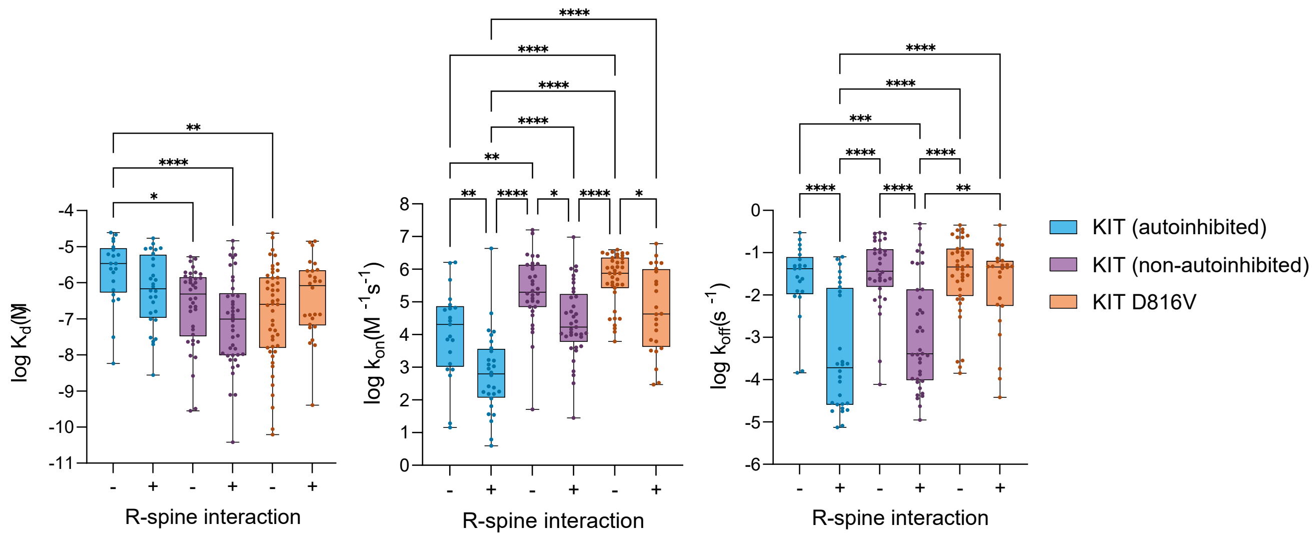


**Figure S5. Distribution of inhibitor binding kinetics and affinity across KIT activation states, stratified by R-spine engagement.** Only compounds within detection limits are included. Statistical significance is indicated as: *, $p < 0.05$; **, $p < 0.01$; ***, $p < 0.001$; ****, $p < 0.0001$.

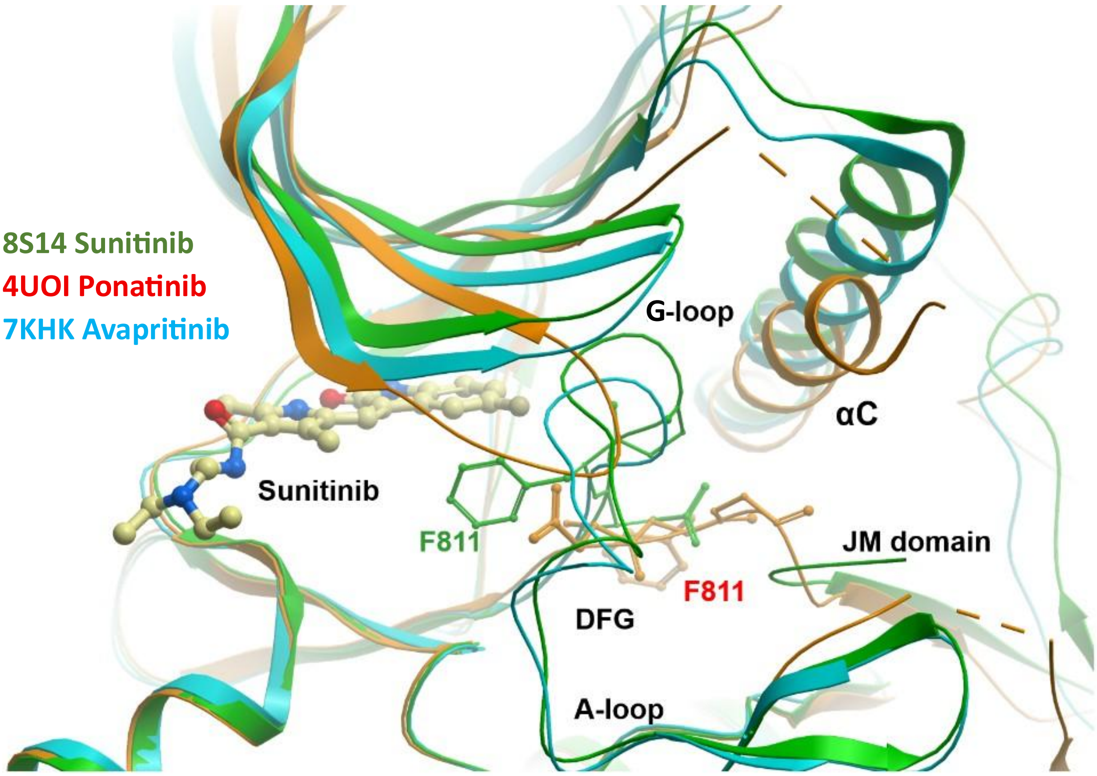


**Figure S6: Superimposition of crystal structures highlighting the plasticity of the KIT kinase domain and diverse conformational states.** Surprisingly Sunitinib (light green; PDB 8S14) stabilizes a DFG-out conformation but does not interact with the allosteric back pocket. Instead, W557 from the JM domain stabilizes an autoinhibited state of the activation segment. Ponatinib (orange, PDB-code 4U0I), the inhibitor is not shown for clarity, binds as expected as a type-II inhibitor inducing a similar conformation of the activation segment as observed in the sunitinib structure. However, the JM does not engage with the kinase domain. αC-helix (αC) is partly unstructured. Avapritinib (olive, PDB-code 7KHK) binds to the active state of KIT. The activation segment assumes a DFG in conformation and it is well ordered. The G-loop bends down towards the ATP binding site and it is rotated outwards. αC-helix is in an “in” position but the N-terminal portion of αC-helix was disordered.

#### SUPPLEMENTARY TABLES

**Table S1. Compounds screened in the HTS kinetic assays.**

| Compound name | Provider | Reference |
|---|---|---|
| 5Z-7-Oxozeaenol | MedChemExpress | HY-12686 |
| Abrocitinib (PF-04965842) | Sigma-Aldrich | SML3475 |
| Acalabrutinib (ACP-196) | MedChemExpress | HY-17600 |
| AEE-788 | MedChemExpress | HY-10045 |
| Afatinib (BIBW2992) dimaleate | SelleckChem | S7810 |
| Afuresertib (GSK2110183) hydrochloride | MedChemExpress | HY-15727A |
| Alectinib (CH5424802) | ApexBio | A8393 |
| Alisertib (MLN8237) | ApexBio | A4110 |
| Alpelisib (BYL719) | SelleckChem | S2814 |
| Alsterpaullone | MedChemExpress | HY-108359 |
| AMG-900 | ApexBio | A4119 |
| Anlotinib (AL3818) | SelleckChem | S8726 |
| Apatinib (YN968D1) | ApexBio | B2303 |
| ARQ-531 | MedChemExpress | HY-112215 |
| Asciminib (ABL001) | SelleckChem | S8555 |
| AST-487 | MedChemExpress | HY-15002 |
| AT-13148 | MedChemExpress | HY-16071 |
| AT-7519 | ApexBio | A5719 |
| AT-9283 | MedChemExpress | HY-50514 |
| Avapritinib (BLU-285) | MedChemExpress | HY-101561 |
| Axitinib (AG 013736) | Sigma-Aldrich | PZ0193 |
| AZ628 | Sigma-Aldrich | SMLO785 |
| AZD-5438 | SelleckChem | S2621 |
| Bafetinib (INNO-406) | MedChemExpress | HY-50868 |
| Barasertib (AZD1152-HQPA/AZD2811) | ApexBio | A4112 |
| Baricitinib (LY3009104, INCB028050) | ApexBio | A4141 |
| BAY-1125976 | MedChemExpress | HY-100018 |
| Binimetinib (MEK162, ARRY-162, ARRY-438162) | ApexBio | A1947 |
| BIRB-796 | SelleckChem | S1574 |
| BLU-9931 | MedChemExpress | HY-12823 |
| BMS-777607 | MedChemExpress | HY-12076 |
| Bosutinib (SKI-606) | Sigma-Aldrich | PZ0192 |
| Brigatinib (AP26113) | ApexBio | A1367 |
| Cabozantinib (XL184) | MedChemExpress | HY13016 |
| Canertinib (CI-1033) dihydrochloride | MedChemExpress | HY-10367 |
| Capivasertib (AZD5363) | MedChemExpress | HY-15431 |
| Capmatinib (INC280) | MedChemExpress | HY-13404 |
| Cediranib (AZD2171) maleate | MedChemExpress | HY-13049 |
| Ceritinib (LDK378) | ApexBio | A3544 |
| CHIR-98014 | MedChemExpress | HY-13076 |
| Cobimetinib (GDC-0973, RG7420) | ApexBio | A3321 |
| Copanlisib (BAY 80-6946) | ApexBio | B2178 |
| Crizotinib (PF-02341066) | ApexBio | B3608 |
| Dabrafenib (GSK2118436) | SelleckChem | S2807 |
| Dacomitinib (PF299804, PF299) | ApexBio | A8319 |
| Danusertib (PHA-739358) | ApexBio | A4116 |

| | | |
|---|---|---|
| Dasatinib (BMS-354825) monohydrate | Santa Cruz | sc-218081 |
| Dinaciclib (SCH 727965) | SelleckChem | S2768 |
| Dovitinib (CHIR-258) | MedChemExpress | HY-50905 |
| Duvelisib (IPI-145, INK1197) | ApexBio | A1720 |
| Elzovantinib (TPX-0022) | MedChemExpress | HY-111787 |
| Encorafenib (LGX818) | ApexBio | B1174 |
| Entrectinib (NMS-E628) | MedChemExpress | HY-12678 |
| Erdafitinib (JNJ-42756493) | MedChemExpress | HY-18709 |
| Erlotinib (CP-358774) hydrochloride | Sigma-Aldrich | CDSO22564 |
| Fasudil (HA-1077) | ApexBio | A5734 |
| Fedratinib (TG101348) | MedChemExpress | HY-10409 |
| Fenebrutinib (GDC-0853) | MedChemExpress | HY-19834 |
| FF-10101 | SelleckChem | S889901 |
| FIIN-2 | MedChemExpress | HY-18602 |
| FIIN-3 | MedChemExpress | HY-18603 |
| Filgotinib (GLPG0634) | MedChemExpress | HY-18300 |
| Fisogatinib (BLU-554) | MedChemExpress | HY-100492 |
| Flavopiridol (HMR-1275) hydrochloride | ApexBio | A8640 |
| Foretinib (XL880) | MedChemExpress | HY-10338 |
| Fostamatinib (R788) | ApexBio | B2284 |
| Futibatinib (TAS-120) | MedChemExpress | HY-100818 |
| Gefitinib (ZD1839) | MedChemExpress | HY-50895 |
| Gilterinib (ASP2215) | SelleckChem | S7754 |
| GSK-1059615 | MedChemExpress | HY-12036 |
| GSK-1070916 | MedChemExpress | HY-70044 |
| GSK-2606414 | ApexBio | A3448 |
| GSK-2656157 | ApexBio | B2175 |
| GSK-690693 | MedChemExpress | HY-10249 |
| GW-5074 | MedChemExpress | HY-10542 |
| H-89 dihydrochoride | MedChemExpress | HY-15979A |
| Ibrutinib (PCI-32765) | ApexBio | A3001 |
| Icotinib (BPI-2009) | ApexBio | A3482 |
| Idelalisib (CAL-101, GS-1101) | ApexBio | A3005 |
| Ilorasertib (ABT-348) hydrochloride | MedChemExpress | HY-16018A |
| Imatinib (STI571) mesylate | Sigma-Aldrich | SML1027 |
| Infigratinib (BGJ-398) | MedChemExpress | HY-13311 |
| Ipatasertib (GDC-0068) dihydrochloride | MedChemExpress | HY-15186A |
| JH-X-119-01 hydrocloride | MedChemExpress | HY-103017 |
| K00546 | MedChemExpress | HY-103647 |
| K02288 | MedChemExpress | HY-12278 |
| Kenpaullone | Selleckchem | S7917 |
| Lapatinib (GW572016) ditosylate | Selleckchem | S1028 |
| Larotrectinib (LOXO-101) | ApexBio | B6176 |
| Lenvatinib (E7080) | ApexBio | A2174 |
| Lestaurtinib (CEP-701) | ApexBio | A4513 |
| Lifirafenib (BGB-283) | MedChemExpress | HY-18957 |
| Linifanib (ABT-869) | ApexBio | A2949 |
| Linsitinib (OSI-906) | ApexBio | A8334 |
| Lorlatinib (PF-6463922) | ApexBio | B4882 |
| LY-3009120 | MedChemExpress | HY-12558 |

| | | |
|---|---|---|
| Masitinib (AB1010) | SelleckChem | S1064 |
| Merestinib (LY2801653) | MedChemExpress | HY-15514 |
| Midostaurin (PKC412) hydrate | Sigma-Aldrich | M1323 |
| Miransertib (ARQ-092) | MedChemExpress | HY-19719 |
| MK-2206 dihydrochoride | MedChemExpress | HY-10358 |
| MK-5108 (VX-689) | ApexBio | A4120 |
| MLN-8054 | ApexBio | A4114 |
| Motesanib (AMG 706) | ApexBio | A3632 |
| Nazartinib (EGF816) | MedChemExpress | HY-12872 |
| Neratinib (HKI-272) | ApexBio | A8322 |
| Netarsudil (AR-13324) | ApexBio | B7807 |
| Nintedanib (BIBF 1120) | ApexBio | A8252 |
| NU-6102 | Santa Cruz | sc-222082 |
| Osimertinib (AZD9291) | ApexBio | B1104 |
| Pacritinib (SB1518) | MedChemExpress | HY-16379 |
| Pazopanib (GW786034) | Sigma-Aldrich | CDS023580 |
| PD-0166285 | MedChemExpress | HY-13925 |
| Pelitinib (EKB-569) | MedChemExpress | HY-32718 |
| Pemigatinib (INCB054828) | MedChemExpress | HY-109099 |
| Pexidartinib (PLX-3397) hydrochloride | MedChemExpress | HY-16749A |
| PF-03814735 | MedChemExpress | HY-14574 |
| PI-103 | ApexBio | A2067 |
| Pictilisib (GDC-0941) | ApexBio | A8210 |
| PLX-4720 | MedChemExpress | HY-51424 |
| Ponatinib (AP24534) | ApexBio | A5467 |
| PP2 | MedChemExpress | HY-13805 |
| Pralsetinib (BLU-667) | MedChemExpress | HY-112301 |
| PRN-1371 | MedChemExpress | HY-101768 |
| Purvalanol A | MedChemExpress | HY-18299A |
| Quizartinib (AC220) | MedChemExpress | HY-13001 |
| Rigosertib (ON-01910) | ApexBio | B1288 |
| Rilzabrutinib (PRN1008) | MedChemExpress | HY-112166 |
| Ripretinib (DCC-2618) | MedChemExpress | HY-112306 |
| RO-31-8220 mesylate | MedChemExpress | HY-13866 |
| Rottlerin | MedChemExpress | HY-18980 |
| Ruxolitinib (INCB018424) | ApexBio | A3012 |
| Sapitinib (AZD8931) | MedChemExpress | HY-13050 |
| SB-202190 | Santa Cruz | sc-202334 |
| Selonsertib (GS-4997) | ApexBio | B7812 |
| Selpercatinib (LOXO-292) | MedChemExpress | HY-114370 |
| Selumetinib (AZD6244) | ApexBio | A8207 |
| Sitravatinib (MGCD516) | MedChemExpress | HY-16961 |
| SNS-032 (BMS-387032) | MedChemExpress | HY-10008 |
| SNS-314 mesylate | ApexBio | A4121 |
| Sonolisib (PX-866) | ApexBio | C3271 |
| SP-600125 | ApexBio | A4604 |
| Staurosporine | ApexBio | A8192 |
| SU-14813 | MedChemExpress | HY-10501 |
| SU-9516 | MedChemExpress | HY-18629 |
| Sunitinib (SU 11248) malate | Sigma-Aldrich | PZ0012 |

| | | |
|---|---|---|
| TAE-684 | ApexBio | A8251 |
| TAK-901 | ApexBio | A4124 |
| Temuterkib (LY3214996) | MedChemExpress | HY-101494 |
| Tepotinib (EMD-1214063) hydrochloride | TargetMol | T9601 |
| THZ1 | ApexBio | A8882 |
| THZ531 | MedChemExpress | HY-103618 |
| Tivozanib (AV-951) | ApexBio | A2251 |
| Tofacitinib citrate | Sigma-Aldrich | PZ0017 |
| Tomivosertib (eFT508) | MedChemExpress | HY-100022 |
| Tozasertib (VX-680, MK-0457) | ApexBio | A4111 |
| Trametinib (GSK1120212) | ApexBio | A3018 |
| Tucatinib (Irbinitinib, ONT-380, ARRY-380) | MedChemExpress | HY-16069 |
| Tyrphostin AG-1478 | MedChemExpress | HY-13524 |
| Upadacitinib (ABT-494) | MedChemExpress | HY-19569 |
| Uprosertib (GSK2141795) | MedChemExpress | HY-15965 |
| Vandetanib (ZD6474) hydrochloride | ApexBio | EDO-015271 |
| Varlitinib (ASLAN001) | MedChemExpress | HY-10530 |
| Vecabrutinib (SNS-062) | MedChemExpress | HY-109078 |
| Vemurafenib (PLX-4032) | SelleckChem | S1267 |
| Verosudil (AR-12286) | MedChemExpress | HY-16758 |
| Vistusertib (AZD2014) | MedChemExpress | HY-15247 |
| Volasetib (BI-6727) | MedChemExpress | HY-12137 |
| VX-702 | MedChemExpress | HY-10401 |
| Zanubrutinib | MedChemExpress | HY-101474A |
| Zimlovisertib (PF-06650833) | MedChemExpress | HY-19836 |
| ZM-336372 | MedChemExpress | HY-13343 |

**Table S2. FDA-approved kinase inhibitors, their main kinase targets and therapeutic indications.**

| Drug name [a] | Trade name | Year approved | Main Targets | Therapeutic indications [b] |
|---|---|---|---|---|
| Abrocitinib | Cibinqo | 2022 | JAK1 | Atopic dermatitis. |
| Acalabrutinib | Calquence | 2017 | BTK | MCL, CLL and SLL. |
| Afatinib | Tovok | 2013 | EGFR, HER2/4 | NSCLC and squamous NSCLC. |
| Alectinib | Alecensa | 2015 | ALK, RET | Third-line Ph+ CML and CML with T315I mutations. |
| Alpelisib | PIQRAY | 2019 | PI3Ka | Breast cancer. |
| Asciminib | Scemblix | 2021 | ABL1 | Ph+ CML. |
| Anlotinib* | FOCUS V | 2018 | KIT, VEGFR1-3 | NSCLC, soft tissue sarcoma, small cell lung cancer and TC. |
| Apatinib* | Aitan | 2014 | VEGFR2 | Gastric cancer. |
| Avapritinib | Ayvakit | 2020 | KIT, KIT D816V, PDGFRa, PDGFRa D842V | Indolent SM, SM, MCL, GISTs with a PDGFRa exon 18 mutation, including D842V. |
| Axitinib | Inlyta | 2012 | KIT, VEGFR1-3 | Advanced RCC. |
| Baricitinib | Olumiant | 2018 | JAK1/2 | Rheumatoid arthritis (RA). |
| Binimetinib | Mektovi | 2018 | MEK1/2 | Melanoma with BRAF V600E or V600K combined with encorafenib. |
| Bosutinib | Bosulif | 2012 | ABL, SRC | Ph+ CML. |
| Brigatinib | Alunbrig | 2017 | ALK, EGFR, FLT3, IGF1R | ALK-positive NSCLC. |
| Cabozantinib | Cometriq & Cabometyx | 2012 | KIT, RET, VEGFR2, MET | Advanced medullary TC, RCC and HCC. |
| Capivasertib | Truqap | 2023 | AKT1/2/3 | HR-positive, HER2-negative breast cancer with PIK3CA/AKT1/PTEN-alterations. |
| Capmatinib | Tabrecta | 2020 | MET | NSCLC with MET exon 14 skipping mutations. |
| Ceritinib | Zykadia | 2014 | ALK, ROS, IGF1R, INSR | ALK-positive NSCLC resistant to crizotinib. |
| Cobimetinib | Cotellic | 2015 | MEK1/2 | BRAF V600E or V600K positive melanomas combined with vemurafenib. |
| Copanlisib | Aliqopa | 2017 | PI3Ka/d | Relapsed follicular lymphoma. |
| Crizotinib | Xalkori | 2011 | ALK, ROS1, MET | ALK- or ROS1-postive NSCLC. |
| Dabrafenib | Tafinlar | 2013 | BRAF, CRAF | BRAF mutation positive melanoma. NSCLC and anaplastic TC with BRAF V600E mutations. |
| Dacomitinib | Visimpro | 2018 | EGFR, HER2/4 | EGFR-mutant NSCLC. |
| Dasatinib | Sprycel | 2006 | KIT, ABL, SRC, LCK, PDGFR | Ph+ CML and ALL. |
| Duvelisib | Copiktra | 2018 | PI3Ka/b/d/g | CLL and SLL. |
| Encorafenib | Braftovi | 2018 | BRAF, BRAF V600E | BRAF V600E or V600K melanoma combined with binimetinib and BRAF V600E colorectal cancer combined with cetuximab. |
| Entrectinib | Rozlytrek | 2019 | TRKA/B/C, ROS1 | Solid tumors with NTRK fusion proteins, ROS1-positive NSCLC. |
| Erdafitinib | Balversa | 2019 | FGFR1-4 | Urothelial bladder cancer. |

| | | | | |
|---|---|---|---|---|
| Erlotinib | Tarceva | 2004 | EGFR | NSCLC and pancreatic cancer. |
| Fedratinib | Inrebic | 2019 | JAK2 | Myelofibrosis. |
| Filgotinib | Jyseleca | 2020 | JAK1/2, TYK2 | RA. |
| Fostamatinib | Tavalisse | 2018 | FLT3, SYK | Chronic immune thrombocytopenia. |
| Futibatinib | Lytgobi | 2022 | FGFR2 | CCA with FGFR2 fusion protein or other rearrangements. |
| Gefitinib | Iressa | 2003 | EGFR | NSCLC with exon 19 deletions or exon 21 substitutions. |
| Gilteritinib | Xospata | 2018 | AXL, FLT3 | FLT3-mutation positive AML. |
| Ibrutinib | Imbruvica | 2013 | BTK | CLL, MCL, marginal zone lymphoma, graft vs. host disease and Waldenström macroglobulinemia. |
| Icotinib* | Conmana | 2011 | EGFR | EGFR-mutation positive advanced NSCLC. |
| Idelalisib | Zydelig | 2014 | PI3Kd | CLL combined with rituximab. |
| Imatinib | Gleevec | 2001 | KIT, ABL, PDGFR | Ph+ CML or ALL, aggressive SM, GISTs, chronic eosinophilic leukemia, hypereosinophilic syndrome, dermatofibrosarcoma protuberans and myelodysplastic or myeloproliferative disease. |
| Infigratinib | Truseltiq | 2021 | FGFR2 | CCA with FGFR2 fusions or other rearrangements. |
| Lapatinib | Tykerb | 2007 | EGFR,HER2 | HER2-positive breast cancer. |
| Larotrectinib | Vitrakvi | 2018 | TRKA/B/C | Solid tumors with NTRK fusion proteins. |
| Lenvatinib | Lenvima | 2015 | KIT, RET, VEGFR1-3 | Differentiated TC, HCC, RCC and endometrial carcinoma. |
| Lorlatinib | Lorbrena | 2018 | ROS1, ALK, ALK L1196M | ALK-positive NSCLC. |
| Midostaurin | Rydapt | 2017 | KIT, FLT3, PKCa/b/g | FLT3 mutation positive AML, SM and mast cell leukemia. |
| Neratinib | Nerlynx | 2017 | EGFR, HER2 | HER2-positive breast cancer. |
| Netarsudil | Rhopressa | 2018 | ROCK1/2 | Glaucoma. |
| Nintedanib | Vargatef | 2014 | KIT, FGFR1-3, VEGFR1-3 | Idiopathic pulmonary fibrosis. |
| Osimertinib | Tagrisso | 2015 | EGFR | NSCLC with exon 19 or 21 substitutions (L858R) or T790M mutations. |
| Pacritinib | Vonjo | 2022 | JAK2 | Myelofibrosis. |
| Pazopanib | Votrient | 2009 | KIT, FGFR1-3, VEGFR1-3 | RCC and soft tissue sarcomas. |
| Pemigatinib | Pemazyre | 2020 | FGFR2 | Advanced CCA with FGFR2 fusions or rearrangements. |
| Pexidartinib | Turalio | 2019 | CSF1R | Tenosynovial giant cell tumors. |
| Ponatinib | Iclusig | 2012 | KIT, ABL1, FGFR, PDGFR | Ph+ CML, including T315I-positive CML and ALL. |
| Pralsetinib | Gavreto | 2020 | RET | RET mutant medullary TC, RET-fusion NSCLC and TC. |
| Quizatinib | Vanflyta | 2023 | FLT3 | FLT3-ITD positive AML combined with cytarabine and daunorubicin. |
| Ripretinib | Qinlock | 2020 | KIT, FLT3, PDGFRa, KDR | Fourth line for GISTs. |
| Ruxolitinib | Jakafi | 2011 | JAK1/2 | Myelofibrosis, polycythemia vera, graft vs. host disease and atopic dermatitis. |

| | | | | |
|---|---|---|---|---|
| Selpercatinib | Retevmo | 2020 | RET | RET mutant medullary TC, RET fusion NSCLC and TC. |
| Selumetinib | Koselugo | 2020 | MEK1/2 | Neurofibromatosis type I. |
| Sunitinib | Sutent | 2006 | KIT, RET VEGFR2, PDGFR | GISTs, RCC and pancreatic neuroendocrine tumors. |
| Tepotinib | Tepmetko | 2021 | MET | MET-mutant NSCL. |
| Tivozanib | Fotvida | 2021 | KIT, VEGFR1-3 | Third line for RCC. |
| Tofacitinib | Tasocitinib | 2012 | JAK1-3 | Ulcerative colitis, RA and psoriatic arthritis. |
| Trametinib | Mekinist | 2013 | MEK1/2 | Melanoma with BRAF V600E or V600K combined with dabrafenib and NSCLC with BRAF V600E combined with dabrafenib. |
| Tucatinib | Tukysa | 2020 | HER2 | HER2-positive breast cancer and colon cancer. |
| Upadacitinib | Rinvoq | 2019 | JAK1 | Atopic dermatitis, second line for RA and psoriatic arthritis. |
| Vandetanib | Zactima | 2011 | VEFGR2 | Medullary TC. |
| Vemurafenib | Zelboraf | 2011 | BRAF | BRAF V600E or V600K positive melanoma combined with cobimetinib and Chester-Erdheim disease. |
| Zanubrutinib | Brukinsa | 2019 | BTK | MCL. |

[a] Kinase inhibitors approved by FDA and NMPA*.

[b] Mantle cell lymphoma (MCL), chronic lymphocytic leukemia (CLL), small lymphocytic lymphoma (SLL), non-small cell lung cancer (NSCLC), Philadelphia chromosome-positive ($Ph^+$), chronic myelogenous leukemia (CML), thyroid cancer (TC), systemic mastocytosis (SM), mast cell leukemia (MCL), gastrointestinal stromal tumors (GISTs), renal cell carcinoma (RCC), rheumatoid arthritis (RA), hepatocellular carcinomas (HCC), acute lymphoblastic leukemia (ALL), acute myeloid leukemia (AML) and cholangiocarcinoma (CCA).

**Table S3. Binding modes of the kinase inhibitors analyzed in this study across relevant proteins.** The binding mode was determined using X-ray crystallography, molecular docking and biochemical assays (BQ) as reported in the literature or KLIFS database. When information for the therapeutically relevant protein was not available, related proteins were used as proxies. For inhibitors whose binding modes were elucidated using crystal structures, the corresponding Protein Data Bank (PDB) entries are provided. In cases where no data was available in the literature, the binding mode was estimated based on the chemical structure and previous structure–activity relationship (SAR) studies or molecular docking. NA marks cases where no information was available. Ref: references.

| Compound | Method | PDB | Protein | Binding mode | Ref. |
|---|---|---|---|---|---|
| 5Z-7-Oxozeaenol | X-ray | 3WZU | MKK7 | VI | 43 |
| Abrocitinib (PF-04965842) | X-ray | 5L2S | CDK6 | $I_{1/2B}$ | 33 |
| Acalabrutinib (ACP-196) | X-ray | 8FD9 | BTK | VI | 33,43 |
| AEE-788 | X-ray | 2ITT | EGFR L858R | I | 34 |
| Afatinib (BIBW2992) dimaleate | X-ray | 4G5J | EGFR | VI | 33,43 |
| Alectinib (CH5424802) | X-ray | 3AOX | ALK | $I_{1/2B}$ | 33,40 |
| Alisertib (MLN8237) | X-ray | 5IA0 | EPHA2 | $I_{1/2B}$ | 40 |
| Alpelisib (BYL719) | X-ray | 7PG6 | PI3Ka | $I_{1/2A}$ | 52 |
| Alsterpaullone | X-ray | 1Q3W | GSK3b | I | 34 |
| AMG-900 | BQ | NA | AURKA | IIA | 44 |
| Anlotinib (AL3818) | NA | NA | VEGFR2 | I | 45 |
| Apatinib (YN968D1) | NA | NA | VEGFR2 | IIA | 44 |
| ARQ-531 | X-ray | 6E4F | BTK | $I_{1/2A}$ | 35 |
| Asciminib (ABL001) | NA | NA | ABL1 | IV | 33 |
| AST-487 | BQ | NA | ABL1 | IIA | 46 |
| AT-13148 | X-ray | 4AXA | PKACA | I | 33 |
| AT-7519 | X-ray | 2VU3 | CDK2 | $I_{1/2B}$ | 34 |
| AT-9283 | X-ray | 2W1G | AURKA | I | 34 |
| Avapritinib (BLU-285) | X-ray | 8PQ9 | KIT | $I_{1/2}$ | 64 |
| Axitinib (AG 013736) | Docking | 1T46 | KIT | IIA | 38 |
| AZ628 | X-ray | 4RZW | BRAF R509H | IIA | 34 |
| AZD-5438 | X-ray | 6GUE | CDK2/CYCA | I | 34 |
| Bafetinib (INNO-406) | X-ray | 2E2B | ABL | IIA | 40 |
| Barasertib (AZD1152- | X-ray | 4C2V | AURKB-INCENP | I | 48 |
| Baricitinib (LY3009104, | X-ray | 6WTO | JAK2 | I | 39 |
| BAY-1125976 | BQ | NA | AKT1/2 | III | 49 |
| Binimetinib (MEK162, ARRY-162, | NA | NA | MEK1/2 | III | 39 |
| BIRB-796 | X-ray | 1KV2 | p38a | IIA | 34 |
| BLU-9931 | X-ray | 4XCU | FGFR4 | VI | 40,43 |
| BMS-777607 | X-ray | 3F82 | MET | IIA | 34 |
| Bosutinib (SKI-606) | X-ray | 3UE4 | ABL1 | IIB | 33 |
| Brigatinib (AP26113) | X-ray | 6MX8 | ALK | $I_{1/2B}$ | 33 |
| Cabozantinib (XL184) | Docking | 1T46 | KIT | IIA | 38,50 |
| Canertinib (CI-1033) dihydrochloride | NA | NA | EGFR | VI | 43 |
| Capivasertib (AZD5363) | X-ray | 4gv1 | AKT1 | I | 33,40 |
| Capmatinib (INC280) | NA | NA | MET | $I_{1/2B}$ | 39 |
| Cediranib (AZD2171) maleate | Chemical | NA | VEGFR2 | I | |
| Ceritinib (LDK378) | X-ray | 4MKC | ALK | $I_{1/2B}$ | 33 |
| Cobimetinib (GDC-0973, RG7420) | X-ray | 4AN2 | MEK1 | III | 33,39 |
| Copanlisib (BAY 80-6946) | X-ray | 5G2N | PIK3CG | $I_{1/2A}$ | 52 |

| | | | | | |
|---|---|---|---|---|---|
| Crizotinib (PF-02341066) | X-ray | 2WGJ | MET | $I_{1/2B}$ | 40 |
| Dabrafenib (GSK2118436) | X-ray | 5CSW | BRAF | $I_{1/2A}$ | 33 |
| Dacomitinib (PF299804, PF299) | X-ray | 4I23 | EGFR | VI | 33,43 |
| Danusertib (PHA-739358) | X-ray | 2J50 | AURKA | IIB | 34 |
| Dasatinib (BMS-354825) | Docking | 1T46 | KIT | IIA | 38 |
| Dinaciclib (SCH 727965) | X-ray | 4KD1 | CDK2 | $I_{1/2B}$ | 40 |
| Dovitinib (CHIR-258) | X-ray | 5AM6 | FGFR1 | $I_{1/2B}$ | 40,51 |
| Duvelisib (IPI-145, INK1197) | NA | NA | NA | I | 52 |
| Elzovantinib (TPX-0022) | X-ray | 8K78 | MET | I | 53 |
| Encorafenib (LGX818) | NA | NA | BRAF | $I_{1/2A}$ | 42 |
| Entrectinib (NMS-E628) | X-ray | 5KVT | TRKA | $I_{1/2B}$ | 33 |
| Erdafitinib (JNJ-42756493) | X-ray | 5EW8 | FGFR1 | $I_{1/2\ A}$ | 40,51 |
| Erlotinib (CP-358774) hydrochloride | X-ray | 1M17 | EGFR | I | 33 |
| Fasudil (HA-1077) | X-ray | 3TKU | MRCKb | I | 40 |
| Fedratinib (TG101348) | X-ray | 6VNE | JAK2 | I | 54 |
| Fenebrutinib (GDC-0853) | X-ray | 5VFI | BTK | $I_{1/2A}$ | 55 |
| FF-10101 | X-ray | 5X02 | FLT3 | VI | 34 |
| FIIN-2 | X-ray | 4QQC | FGFR4 | VI | 43 |
| FIIN-3 | X-ray | 4R6V | FGFR4 V550L | VI | 43 |
| Filgotinib (GLPG0634) | X-ray | 4P7E | JAK1 | I | 34 |
| Fisogatinib (BLU-554) | X-ray | 6NVK | FGFR4 | VI | 43 |
| Flavopiridol (HMR-1275) | X-ray | 3BLR | CDK9/CycT1 | I | 34 |
| Foretinib (XL880) | X-ray | 3LQ8 | MET | IIA | 40,44 |
| Fostamatinib (R788) | X-ray | 3FQS | SYK | I | 33,40 |
| Futibatinib (TAS-120) | X-ray | 6MW | FGFR1 | VI | 33,43 |
| Gefitinib (ZD1839) | X-ray | 2ITY | EGFR | I | 33 |
| Gilterinib (ASP2215) | X-ray | 6jqr | FLT3 | IIB | 33 |
| GSK-1059615 | Chemical | NA | PI3K | I | |
| GSK-1070916 | BQ | NA | AURKB/C | IIB | 41 |
| GSK-2606414 | X-ray | 4G31 | PERK | $I_{1/2A}$ | 34 |
| GSK-2656157 | X-ray | 4M7I | PERK | $I_{1/2A}$ | 34 |
| GSK-690693 | X-ray | 3D0E | AKT2 | I | 40 |
| GW-5074 | NA | NA | CRAF | I | 42 |
| H-89 dihydrochoride | X-ray | 3FMD | HASPIN | I | 40 |
| Ibrutinib (PCI-32765) | X-ray | 5P9J | BTK | VI | 40,43 |
| Icotinib (BPI-2009) | NA | NA | EGFR | I | 56 |
| Idelalisib (CAL-101, GS-1101) | X-ray | 4XE0 | PI3Kd | I | 52 |
| Ilorasertib (ABT-348) hydrochloride | Docking | NA | RET | II | 57 |
| Imatinib (STI571) mesylate | X-ray | 1T46 | KIT | IIA | 40,63 |
| Infigratinib (BGJ-398) | X-ray | 3TT0 | FGFR1 | $I_{1/2B}$ | 33 |
| Ipatasertib (GDC-0068) | X-ray | 4EKL | AKT1 | I | 40 |
| JH-X-119-01 hydrocloride | NA | NA | IRAK1 | VI | 43 |
| K00546 | X-ray | 2WU6 | CLK3 | I | 40 |
| K02288 | X-ray | 6EIX | ACVR1 Q207E | $I_{1/2B}$ | 34 |
| Lapatinib (GW572016) ditosylate | X-ray | 1XKK | EGFR | $I_{1/2A}$ | 33 |
| Larotrectinib (LOXO-101) | NA | NA | TRKA/B/C | I | 33 |
| Lenvatinib (E7080) | Docking | 1T46 | KIT | IIA | 38 |
| Lestaurtinib (CEP-701) | X-ray | 8BPW | JAK2 | I | 34 |
| Lifirafenib (BGB-283) | X-ray | 4R5Y | BRAF | IIA | 34 |
| Linifanib (ABT-869) | NA | NA | VEGFR2 | IIA | 59 |

| | | | | | |
|---|---|---|---|---|---|
| Linsitinib (OSI-906) | Chemical | NA | IGF1, INSR | I | |
| Lorlatinib (PF-6463922) | X-ray | 4CLI | ALK | I | 33 |
| LY-3009120 | X-ray | 5C9C | BRAF V600E | IIA | 34 |
| Masitinib (AB1010) | NA | NA | KIT | IIA | 44 |
| Merestinib (LY2801653) | X-ray | 4EEV | MET | IIA | 34 |
| Midostaurin (PKC412) hydrate | Docking | 1PKG | KIT | I | 38 |
| Miransertib (ARQ-092) | X-ray | 5KCV | AKT1 | III | 34 |
| MK-2206 dihydrochoride | NA | NA | AKT1/2/3 | III | 49 |
| MK-5108 (VX-689) | X-ray | 5EW9 | AURKA | $I_{1/2B}$ | 40 |
| MLN-8054 | X-ray | 2WTV | AURKA | IIB | 34 |
| Motesanib (AMG 706) | X-ray | 3EFL | VGFR2 | IIA | 40 |
| Nazartinib (EGF816) | X-ray | 5FEQ | EGFR | VI | 43 |
| Neratinib (HKI-272) | X-ray | 3W2Q | EGFR | VI | 40,43 |
| Netarsudil (AR-13324) | NA | NA | NA | I | 39 |
| Nintedanib (BIBF 1120) | X-ray | 3C7Q | VEGFR2 | IIB | 33 |
| NU-6102 | X-ray | 5LQF | CDK1/CycB | I | 40 |
| Osimertinib (AZD9291) | X-ray | 6JXT | EGFR | VI | 33,43 |
| Pacritinib (SB1518) | X-ray | 8BPV | JAK2 | I | 60 |
| Pazopanib (GW786034) | NA | NA | NA | I | 39 |
| PD-0166285 | Chemical | NA | WEE1 | I | |
| Pelitinib (EKB-569) | NA | NA | EGFR | VI | 43 |
| Pemigatinib (INCB054828) | X-ray | 7WCL | FGFR1 | I | 39 |
| Pexidartinib (PLX-3397) | X-ray | 7KHG | KIT | IIA | 34 |
| PF-03814735 | X-ray | 5VD2 | WEE1 | I | 34 |
| PI-103 | X-ray | 4L23 | PI3Ka | I | 34 |
| Pictilisib (GDC-0941) | X-ray | 2WXP | PI3Kd | I | 34 |
| PLX-4720 | X-ray | 3C4C | BRAF | $I_{1/2A}$ | 34 |
| Ponatinib (AP24534) | X-ray | 4U0I | KIT | IIA | 38 |
| PP2 | X-ray | 3SXS | BMX | II | 40 |
| Pralsetinib (BLU-667) | X-ray | 7JU5 | RET | I | 34 |
| PRN-1371 | X-ray | 7F3M | FGFR4 | VI | 43 |
| Purvalanol A | X-ray | 1YOM | SRC | IIB | 34 |
| Quizartinib (AC220) | X-ray | 4xuf | FLT3 | IIA | 33 |
| Rigosertib (ON-01910) | NA | NA | PLK1 | III | 62 |
| Rilzabrutinib (PRN1008) | X-ray | 7L5P | BTK | VI | 34 |
| Ripretinib (DCC-2618) | X-ray | 6MOB | KIT | IIA | 33 |
| RO-31-8220 mesylate | X-ray | 4OTH | PRK1 | $I_{1/2}$ | 40 |
| Rottlerin | Chemical | NA | PKC | I | |
| Ruxolitinib (INCB018424) | X-ray | 6VGL | JAK2 | I | 33 |
| Sapitinib (AZD8931) | Docking | NA | EGFR | I | 65 |
| SB-202190 | Chemical | NA | p38a | I | |
| Selonsertib (GS-4997) | X-ray | 6OYT | ASK1 | I | 34 |
| Selpercatinib (LOXO-292) | X-ray | 7JU6 | RET | $I_{1/2B}$ | 33 |
| Selumetinib (AZD6244) | X-ray | 4U7Z | MEK1 | III | 33 |
| Sitravatinib (MGCD516) | NA | NA | NA | II | 61 |
| SNS-032 (BMS-387032) | X-ray | 5D1J | CDK2 | $I_{1/2B}$ | 34 |
| SNS-314 Mesylate | X-ray | 3D15 | AURKA | I | 41 |
| Sonolisib (PX-866) | NA | NA | PI3K | VI | 43 |
| SP-600125 | X-ray | 2ZMD | ITK T686A | I | 34 |
| Staurosporine | X-ray | 4QMY | STK24 | I | 40 |

| | | | | | |
|---|---|---|---|---|---|
| SU-14813 | Chemical | NA | VEGFR | II | |
| SU-9516 | X-ray | 6GUC | CDK2/CycA | I | 34 |
| Sunitinib (SU 11248) malate | X-ray | 3G0E | KIT | IIB | 38 |
| TAE-684 | X-ray | 2XB7 | ALK | I | 34 |
| TAK-901 | BQ | NA | NA | I | 41 |
| Temuterkib (LY3214996) | X-ray | 6RQ4 | ERK2 | I | 34 |
| Tepotinib (EMD-1214063) | X-ray | 4R1V | MET | $I_{1/2B}$ | 33 |
| THZ1 | X-ray | 6XD3 | CDK7 | VI | 43 |
| THZ531 | X-ray | 7NXJ | CDK13 | VI | 43 |
| Tivozanib (AV-951) | X-ray | 4ASE | VEGFR2 | IIA | 33 |
| Tofacitinib citrate | X-ray | 3EYG | JAK1 | $I_{1/2B}$ | 33,40 |
| Tomivosertib (eFT508) | X-ray | 6CK6 | MNK2 D228G | I | 34 |
| Tozasertib (VX-680, MK-0457) | X-ray | 3E5A | AURKA-TPX2 | I | 34 |
| Trametinib (GSK1120212) | NA | NA | NA | III | 33,39 |
| Upadacitinib (ABT-494) | NA | NA | NA | I | |
| Vandetanib ZD6474) hydrochloride | X-ray | 2IVU | RET | I | |
| Vemurafenib (PLX-4032) | X-ray | 4RZV | BRAF | I | 39 |
| Verosudil (AR-12286) | Chemical | NA | ROCK1 | II | |
| Volasetib (BI-6727) | X-ray | 3FC2 | PLK1 | I | 33 |
| VX-702 | Chemical | NA | p38a | $I_{1/2A}$ | 33 |
| Zanubrutinib | X-ray | 6J6M | BTK | IIA | |
| Zimlovisertib (PF-06650833) | X-ray | 5UIU | IRAK4 | I | 34 |
| ZM-336372 | NA | NA | BRAF | I | |
